# Hippocampus single-nucleus transcriptomics reveals coordinated regulation of social and spatial representation development by perinatal SERT expression in CA3 pyramidal neurons

**DOI:** 10.64898/2026.05.11.724399

**Authors:** Wenna Chen, Roberto De Gregorio, Maider Astorkia, Ji Ying Sze, Deyou Zheng

## Abstract

The hippocampal formation (HPF) provides neural substrates integrating disparate sensory cues into episodic memories and coherent action. Whereas HPF structures are formed by birth, the functional circuits evolve over postnatal development. Our previous studies showed that transient perinatal expression of the serotonin (5-HT) transporter SERT/*Slc6a4* in CA3 pyramidal neurons, which do not synthesize 5-HT but take up extracellular 5-HT thus termed “5-HT-absorbing neurons”, exerts sex-biased effects on long-term activity-dependent HPF synaptic plasticity and behavior in mice. This study investigates SERT impact on circuit development, through single-nucleus transcriptomics of postnatal HPF from CA3-pyramidal neuron SERT knockout (*SERT^PyramidΔ^*) mice. We demonstrate that *SERT^PyramidΔ^* mice preserve cell identities across the HPF but alter gene expression in specific neuronal types in a sex-biased manner. We observed *SERT^PyramidΔ^* male-biased upregulation of genes preferentially in glutamatergic neurons, particularly affecting the CA2 and parasubiculum (PaS) when they develop social novelty and spatial representations, respectively. In both the CA2 and PaS, altered genes center on two categories –– modulators of gene expression patterning including chromatin plasticity, RNA processing and ubiquitin-dependent protein degradation, and aspects of synaptic transmission. >20% of the dysregulated genes in the CA2 and PaS are associated with Autism and engaged in cell-type distinct functional networks, showing CA3 SERT regulation of ASD-vulnerable genes in intersecting biological processes in specific neurons during social and spatial circuits development. The data, available at https://scviewer.shinyapps.io/hippocampus_sertKO, provide an entry map for further deducing anatomical neuronal origin and the molecular and cellular pathways impaired by 5-HT dysfunction during HPF circuits development leading to lifetime cognitive deficits.

## INTRODUCTION

The hippocampal formation (HPF) mediates declarative learning by integrating sensory information about the environment and physiological demands into episodic memories, and underscores many neurological disorders (Milner et al., 1998; O’Keefe and Nadel, 1978). The HPF comprises four regions –– the hippocampus proper of three *Cornu Ammonis* fields (CA1, CA2, CA3), the dentate gyrus (DG), the subicular complex, and the entorhinal cortex (EC) (Amaral and Witter, 1989; van Strien et al., 2009). Each HPF region embodies unique neuronal cells with distinct gene expression patterns and synaptic connectivity (Cembrowski et al., 2016; Ding et al., 2020; Lavenex and Banta Lavenex, 2013). To understand HPF contributions to cognitive and behavioral deficits and to develop targeted therapies, we need to determine the cellular origin –– where in the HPF, in which cell types and when during the development ––that is impaired by particular genetic variants and environmental insults causing observed phenotypes.

The gross HPF anatomy and the neuronal repertoire are established by birth; the functional circuits, however, evolve over postnatal development (Lavenex and Banta Lavenex, 2013; Leinekugel, 2003; Nadel, 2022). In mice, the first two postnatal weeks of neural circuit development is a “critical period” of plasticity –– changes in this period may permanently alter circuit configuration (Hensch, 2005; Marin, 2016; McHail and Dumas, 2015). Great strides have been made in identifying the role of particular HPF regions in behavioral circuits (Dalton and Maguire, 2017; Hitti and Siegelbaum, 2014). As accurate episodic memories and coherent behavior require precise coordination of distinct neural circuits across the HPF, identification of regulatory mechanisms that coordinate the development of functional related neural circuits in the HPF during the critical period is a crucial next step towards understanding behavior in health and disease.

The HPF is sexually dimorphic at the level of gene expression, cellular dynamics, and synapse organization (Bundy et al., 2017; Lenz et al., 2012; Premachandran et al., 2020). Most neuropsychiatric disorders display sex-biased phenotypes (Labonté et al., 2017; Satterthwaite et al., 2015). For instance, males have 2 - 4 times higher risks than females to manifest characteristic behavioral deficits of autism spectrum disorders (ASD) (Global Burden of Disease Study Autism Spectrum, 2025). Analyzing postmortem cerebral cortex transcriptomics of ASD subjects identified small magnitude, male-biased gene upregulation (Werling et al., 2016). Further, an ASD-associated allele of the chromatin-remodeling factor *Chd8* confers male-biased upregulation as well as male-female opposite changes in gene expression and sex-specific aberrant HPF synaptic activities and behavior in mice (Jung et al., 2018). Moreover, male but not female carriers of genetic variants in the synapse scaffolding protein Shank1 and ryanodine receptor 2 (RyR2) manifest ASD (Lu and Cantor, 2012; Sato et al., 2012). Key question remains: when sex-biased circuit derailments in disease brain may arise?

In the brain, the serotonin uptake transporter (SERT/*Slc6a4*) is the primary mechanism for clearing extracellular serotonin (5-HT), thereby defining 5-HT availability to the receptors (Hahn and Blakely, 2007). The effects of 5-HT dysfunction on psychiatric disorders are sex biased (Frey et al., 2010; Parsey et al., 2002; Seney et al., 2018). Importantly, 5-HT actions in developing and adult brain differ. Whereas in adult brain 5-HT is a neurotransmitter released at the synapses, in developing brain 5-HT is mainly a trophic factor as raphe serotonergic neurons release 5-HT throughout the brain before synapse formation (Azmitia, 1999; Gaspar et al., 2003). G. Buznikov and J. Lauder demonstrated that trophic 5-HT is an ancient morphogen influencing spatial-temporal organization of ontogenesis (Buznikov et al., 2001; Colgan et al., 2009). Pioneering work by P. Gaspar and colleagues identified in multiple organisms that in the developing brain SERT is expressed in defined sets of non-serotonergic neurons, in addition to raphe serotonergic neurons (Lebrand et al., 1996; Lebrand et al., 1998). Several groups including ourselves have confirmed in rodents that SERT is expressed from embryonic day (E) 17 to postnatal day (P) 10 specifically in subsets of thalamocortical projection neurons and pyramidal neurons located in the medial prefrontal cortex (mPFC) and hippocampal CA3 (Chen et al., 2015; De Gregorio et al., 2022; Hansson et al., 1998; Soiza-Reilly et al., 2019). These SERT-expressing neurons –– termed “5-HT-absorbing neurons” –– take up then degrade trophic 5-HT from the extracellular space thereby defining 5-HT levels during the critical period of postnatal neural circuit development (Chen et al., 2015; Lebrand et al., 1996).

By generating mice with SERT expression ablated in specific neurons, we and others have demonstrated that SERT expression in 5-HT-absorbing neurons is not required for achieving the identity of themselves or their target cells, but is essential for elaborating normal synaptic structure at the target brain regions (Chen et al., 2015; De Gregorio et al., 2020). Specifically, ablating SERT in the 5-HT-absorbing pyramidal neurons (*SERT^PyramidΔ^*) resulted in increased functional synapses at mPFC and CA3 target regions in the adult brain (De Gregorio et al., 2022; Soiza-Reilly et al., 2019) and sex-biased impairments in long-term activity-dependent hippocampal synaptic plasticity and cognitive behaviors (De Gregorio et al., 2022).

The current study aims to link aberrant synaptic structure and cognitive behaviors to the critical period of circuit assembly at molecular and cellular levels across the HPF in *SERT^PyramidΔ^* mice. We employed single-nucleus RNA-sequencing (snRNA-seq) to profiling gene expression of the dorsal HPF (dHPF) in male and female *SERT^PyramidΔ^ vs*. control littermates aged P16, at the end of the critical developmental period while cognitive behaviors emerging. In line with the hypothesis that SERT expression in 5-HT-absorbing neurons serves to define spatiotemporal trophic 5-HT signaling (Gaspar et al., 2003; Jafari et al., 2011), we found *SERT^PyramidΔ^* effects not conformed to CA3 pyramidal neuron synaptic partners, but selectively on neuronal types representing social and spatial perceptions via sex biased mechanisms. *SERT^PyramidΔ^* males preferentially altered gene expression in glutamatergic neurons in the CA2 and parasubiculum (PaS) when they develop social and spatial representations, respectively. In both the CA2 and PaS, *SERT^PyramidΔ^*-affected genes coalesced in cellular processes for chromatin plasticity, RNA processing, ubiquitin (Ubi)-dependent protein degradation and aspects of synaptic transmission, but the genes affected in these processes were largely neuron-type specific. Congruent with the notion that a myriad of genetic risk factors may converge on intersecting biological processes in critical developmental timing to contribute to common phenotypes (Parikshak et al., 2013; Willsey et al., 2013), >20% of *SERT^PyramidΔ^*-affected genes in the CA2 and PaS, engaged in cell-type distinct functional networks, are associated with ASD. These data, available at https://scviewer.shinyapps.io/hippocampus_sertKO, shed new light into the role of SERT and trophic 5-HT in shaping normal HPF functional circuits development and disease.

## RESULTS

### Generation of postnatal HPF single-nucleus transcriptomics of *SERT^PyramidD^* mice

Previous studies have demonstrated that in the mouse brain SERT is transiently expressed specifically in subsets of pyramidal neurons located in the mPFC and CA3 (De Gregorio et al., 2022; Soiza-Reilly et al., 2019). In both males and females, SERT-expressing CA3 pyramidal neurons emerge at E17.5, progressively growing their axonal projections to innervate the stratum oriens, stratum radiatum and DG at both the ipsilateral and contralateral HPF during the first postnatal week (Figures 1A and 1B), and the SERT expression is terminated by P10 (De Gregorio et al., 2022; Lebrand et al., 1998). The SERT-expressing mPFC pyramidal neurons are located in layers 5 and 6, projecting to a wide range of cortical and subcortical regions but not the HPF (Figure 1B) (De Gregorio et al., 2022; Soiza-Reilly et al., 2019). *SERT^PyramidΔ^* mice specifically ablate SERT expression in the pyramidal neurons (De Gregorio et al., 2022). As the first step towards delineating the lasting effects of spatiotemporal SERT expression on gene expression in distinct cell types in the HPF, we performed snRNA-seq of the HPF from male and female *SERT^PyramidΔ^* and control *SERT^fl/fl^* littermates at P16, the timing when pup’s eyelid is parting and higher cognitive behaviors begin to emerge (Figure 1C).

**Figure 1.**
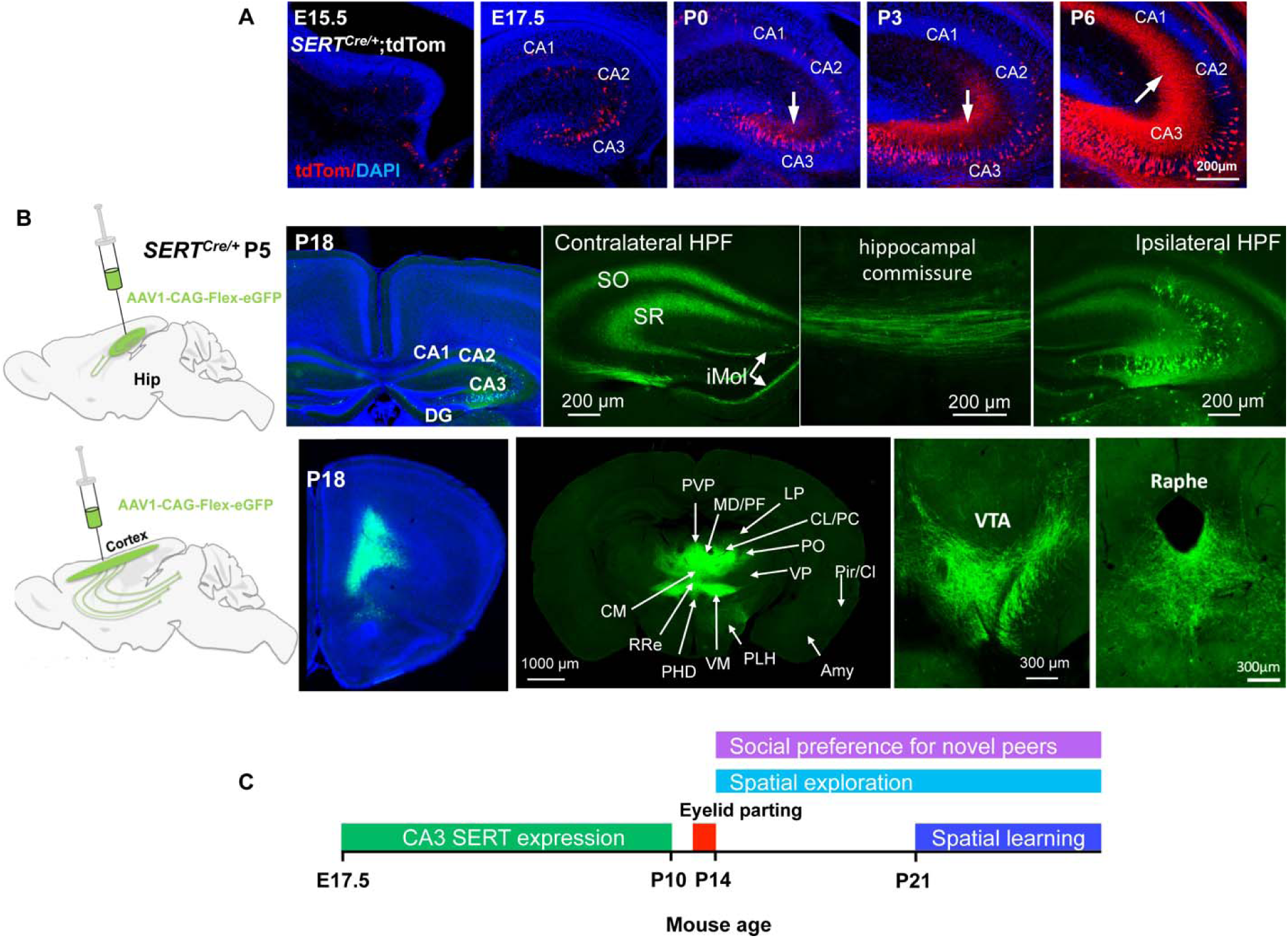
SERT is expressed in CA3 pyramidal neurons specifically during HPF functional circuit development. **A.** *SERT^Cre/+-^*dependent tdTom-expressing neurons arise at E17.5, progressively increasing in number and extending axonal projections (indicated by arrows) in the HPF (outlined by DAPI counterstain in blue) during the first postnatal week. **B.** SERT-expressing pyramidal neuron axonal projection network revealed by AAV-mediated anterograde tracing. **Top panels:** left, schematic of unilateral stereotaxic injection of AAV1 expressing Cre-dependent GFP (AAV1.CAG.Flex.eGFP.WPRE.bGH) into the HPF of P5 *SERT^Cre/+^*pups, second to the left, a photomicrograph showing GFP+ neurons at the injection site when the mouse aged P18, with the HPF outlined by DAPI counterstain (blue), and right 3 photomicrographs showing GFP+ axonal projections to the stratum oriens (SO), stratum radiatum (SR) and DG inner-molecular layer (iMol) both at the ipsilateral HPF and via the commissural pathway to the contralateral HPF. **Bottom panels**: left, schematic of unilateral stereotaxic injection of Cre-dependent GFP-expressing AAV1 into the mPFC of P5 *SERT^Cre/+^* pups, second to the left, a photomicrograph showing GFP+ neurons at the injection site when the mouse aged P18, with the cortex outlined by DAPI counterstain (blue), and right three photomicrographs showing GFP+ axonal projections to many cortical and subcortical regions but not HPF. Amy, Amygdala; CM, central medial thalamic nucleus; CL, centrolateral thalamic nucleus; PC, paracentral thalamic nucleus; LP, lateral posterior thalamic nucleus; MD, mediodorsal thalamic nucleus; PF, parafascicular thalamic nucleus; PHD, posterior hypothalamic area; Pir, piriform cortex; Cl, Claustrum; PLH, peduncular part of lateral hypothalamus; PO, posterior thalamic nuclear group; PVP, paraventricular thalamic nucleus; RRe, retrouniens area; VM, ventromedial thalamic nucleus; VP, ventral posterior thalamic nucleus; VTA, ventral tegmental area. The images in A and B are adapted with permission from our published work (De Gregorio et al., 2022) and included here simply to help visualization of the structural content of the current study. **C.** Timing of SERT expression in CA3 pyramidal neurons in relation to eyelid parting and the emergence of social novelty preference and spatial exploration behavior.

We implemented several strategies to enhance transcriptome resolution and specificity (Figure 2A). First, we dissected the HPF free from the EC and surrounding brain tissues. Second, since the dorsal (d) and ventral (v) HPF differ in cell type compositions, neuronal properties and gene expression patterns (Amaral and Witter, 1989; Cembrowski et al., 2016; Yao et al., 2021; Zeisel et al., 2015) and dHPF preferentially mediates cognitive functions (Fanselow and Dong, 2010; Kheirbek and Hen, 2011), we elected to perform RNA-seq of nuclei isolated from the dHPF. Third, to minimize inter-animal biological variations each sequencing sample contained pooled nuclei of sex/genotype-matched dHPF from multiple independent cohorts (n = 5 male mice/genotype, 6 female mice/genotype). An aliquot of the nuclei was used to generate snRNA-seq libraries using 10x Genomics Chromium v3.1, targeting 350 million sequencing reads per sample.

**Figure 2.**
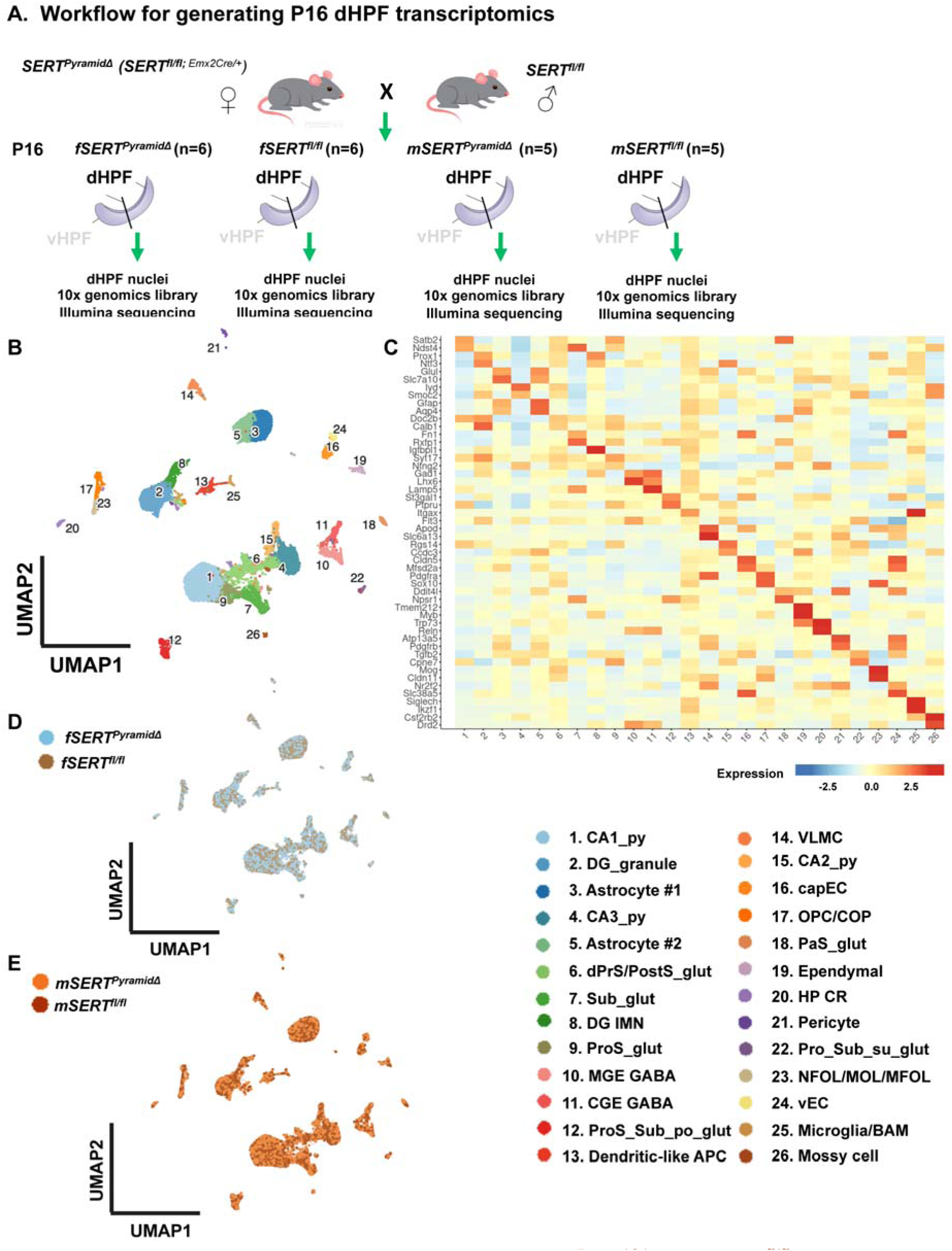
Transcriptomic cell types in dHPF of *SERT^PyramidΔ^* and *SERT^fl/fl^* mice at P16. **A.** Workflow for generating single-nucleus transcriptomics of the dHPF from *SERT**^Pyramid^**^Δ^* and *SERT^fl/fl^* mice aged P16. n = the number of mice used in indicated sample. **B.** UMAP representation of all single nucleus transcriptomes, colored by cell class. APC, antigen-presenting cell; BAM, border-associated macrophage; capEC, capillary endothelial cell; CGE, caudal ganglionic eminence; COP, committed oligodendrocyte precursor; dPrS/PostS dorsal presubiculum/postsubiculum; glut, glutamatergic neuron; HP CR, hippocampal Cajal-Retzius cell; IMN, immature neuron; MGE, medial ganglionic eminence; MOL, mature oligodendrocyte; MFOL, myelin-formed oligodendrocyte; NFOL, newly formed oligodendrocyte; OPC, oligodendrocyte progenitor cell; PaS, parasubiculum; po, polymorphic cell layer; ProS, prosubiculum; py, pyramidal neuron; su, superficial layer; Sub, subiculum proper; vEC, venous endothelial cells, VLMC, vascular leptomeningeal cell. **C.** Heatmap showing the expression of selected cell-type markers in transcriptomic clusters. Colors indicate relative cluster mean expression levels. **D – E.** UMAP representations showing all single nucleus transcriptomes colored by sex and genotype, female *SERT^PyramidΔ^ vs.* female *SERT^fl/fl^* samples (**D**), and male *SERT^PyramidΔ^ vs.* male *SERT^fl/fl^*samples (**E**).

Following strict quality control (QC) filtering, we retained 52,047 high-quality transcriptomes representing ∼26,000 genes in male and female *SERT^PyramidΔ^* and *SERT^fl/fl^* dHPF (Figure S1). The datasets from all the samples were integrated and clustered for differential analysis using our recently developed pipeline (Ferrena et al., 2024), and are available at https://scviewer.shinyapps.io/hippocampus_sertKO.

### *SERT^PyramidΔ^* mice preserve normal cell type identities across HPF

We first assessed cell populations in the male and female *SERT^PyramidΔ^ vs*. *SERT^fl/fl^* HPF transcriptomes using unbiased clustering. This identified 26 clusters as visualized by uniform manifold approximation and projection (UMAP) (Figure 2B). We next sought to determine cluster cell-type identities, using consensus gene markers from HPF-specific transcriptomics, mouse brain spatial transcriptomics and Allen Brain Atlas RNA in situ hybridization data (Cembrowski et al., 2016; Ding et al., 2020; Lein et al., 2007; Yao et al., 2023; Yao et al., 2021; Zhang et al., 2023). We identified eleven of the clusters each representing glutamatergic neurons in anatomically distinct HPF regions –– pyramidal neurons of CA1, CA2 and CA3, DG mossy cells, DG granule cells, and six populations in the subicular complex (Figures 2B and 2C). We validated that the transcriptomes primarily represent dHPF cell types, by examining gene markers selective for dorsal or ventral CA1, CA3 pyramidal neurons and DG granules (Cembrowski et al., 2016; Dong et al., 2009; Lein et al., 2007; Yao et al., 2021). In each of the cell-type clusters, we observed robust expression of the dorsal cell markers, contrasting to minimal expression of the ventral markers (Figure S2A).

We identified two clusters representing GABAergic neurons of different developmental origins: the medial ganglionic eminence (MGE) and caudal ganglionic eminence (CGE) (Figures 2B and 2C). MGE and CGE each give rise to multiple functionally distinct GABAergic neuron subtypes that express characteristic neuropeptides and receptors (Paul et al., 2017; Wamsley and Fishell, 2017). Accordingly, we detected the somatostatin gene *Sst* and parvalbumin gene *Pvalb* expression in subpopulations in the MGE GABA cluster, and the vasointestinal peptide gene *Vip*, ionotropic 5-HT receptor gene *Htr3a* and extracellular matrix glycoprotein Reelin gene *Reln* in the CGE GABA cluster (Figure S2B).

We identified two clusters representing HPF-specific immature neurons: glutamatergic DG immature neurons (DG-IMN) and hippocampal Cajal-Retzius (HP CR) cells (Figures 2B, 2C and S2C). ∼85% of DG granule cells in the rodent brain are born postnatally, with the proliferation peaking around the end of the first postnatal week (Altman and Das, 1966; Bayer, 1980; Schlessinger et al., 1975). Unlike embryonically generated granule cells, which migrate to the DG from the ventricular zone, postnatally-generated granule cells are born locally in the proliferative zone in the DG hilar region (Bayer, 1980; Rakic and Nowakowski, 1981; Schlessinger et al., 1975). The DG-IMN cluster displayed minimal expression of early neuroblast markers (*Neurod4, Mfap4, Eomes, Zic5*), but robust late neuroblast/immature DG granule cell markers (*Igfbpl1, Draxin, Fam163a, Rarb*) (Figure S2C), indicating the majority of which in the process to be incorporated into DG circuits during the functional maturation.

In addition to the robust expression of CR cell hallmarks *Trp73*, *Reln*, HPF-specific CR cell markers *Ebf3*, *Ndnf*, we observed *Slc17a6* (vesicular glutamate transporter Vglut2) expression in a subpopulation of the CR cell cluster (Figure S2C). While CR cells are mostly transient neurons secreting Reelin to guide migrating neurons and axonal wiring during HPF morphogenesis, a small amount of CR cells persist in the HPF using Vglut2 for glutamate transmission and influencing HPF synaptic circuit maturation (Anstotz et al., 2016; Anstotz et al., 2022; Quattrocolo and Maccaferri, 2014).

We identified eleven clusters representing four classes of non-neuronal cells (Figures 2B and 2C). (1) Vascular cells: vascular leptomeningeal cells/VLMC, capillary endothelial cells/capEC, venous endothelial cells/vEC and pericytes. (2) Glial cells: Astrocyte #1, Astrocyte #2, populations of oligodendrocyte precursor cells/OPC and committed oligodendrocyte precursors/COP, and oligodendrocytes at various maturation stages (newly formed oligodendrocytes/NFOL, mature oligodendrocytes/MOL, myelin-formed oligodendrocytes/MFOL). (3) Immune cells: dendritic-like antigen-presenting cell/APC populations, microglia and border-associated macrophages/BAM. (4) Ependymal cells. The two astrocyte clusters shared in common many canonical astrocyte markers, including transcription factors (e.g. *Gli2*, *Gli3*, *Rfx4*) and functional components (e.g. *Glul*, *Aldh1l1*), but differed in that Astrocyte #1 showed low *Gfap* and *Aqp4* but robust *Slc7a10* expression, whereas Astrocyte #2 displayed higher *Gfap* and *Aqp4* expression (Figure S2D). *Slc7a10* encodes an amino acid transporter that in the HPF is primarily expressed in astrocytes to mediate sodium-independent glycine efflux for the homeostasis of excitatory and inhibitory neurotransmission (Batiuk et al., 2020; Ehmsen et al., 2016). Therefore, Astrocyte clusters #1 and #2 likely represent functionally distinct astrocyte subpopulations. Likewise, the two clusters of immune cells displayed many markers in common (Figure S2E). The dendritic-like APC cluster was heterogeneous, distinguished by the low expression of the microglia markers *Tmem119* and *P2ry12* relative to the major histocompatibility complex markers *Itgax/Cd11c, Flt3, Cd74, Ciita* and *Cd24a* (Figure S2E)(Boss, 1997; Bridlance and Thion, 2023; Bulloch et al., 2008; Rovira et al., 2025).

Together, our snRNA-seq transcriptomics covered expected neuronal and non-neuronal cell types across the dHPF. The resolved cell subtypes indicate high quality and high resolution of the datasets. The *SERT^PyramidΔ^* transcriptomes showed the same cell populations as the controls (Figures 2D and 2E), supporting previous findings that SERT expression in the 5-HT-absorbing pyramidal neurons is not required for gross cell-fate specification of CA3 pyramidal neurons or their targets in the HPF (De Gregorio et al., 2022).

### Sex-biased variations in HPF cell type composition

Across the four samples, dHPF transcriptomes contained 62% - 67% neurons, 26% - 32% non-neuronal cells and 3% - 7% immature neurons (Figures 3A and 3B; Table S1). Among the neurons, 89% - 92% were glutamatergic neurons and 8% - 11% GABAergic neurons (Figures 3A and 3C; Table S1), in line with ∼90% glutamatergic neurons *vs.* ∼10% GABAergic neurons observed in adult mouse HPF single-cell transcriptomics (Zhang et al., 2023).

**Figure 3.**
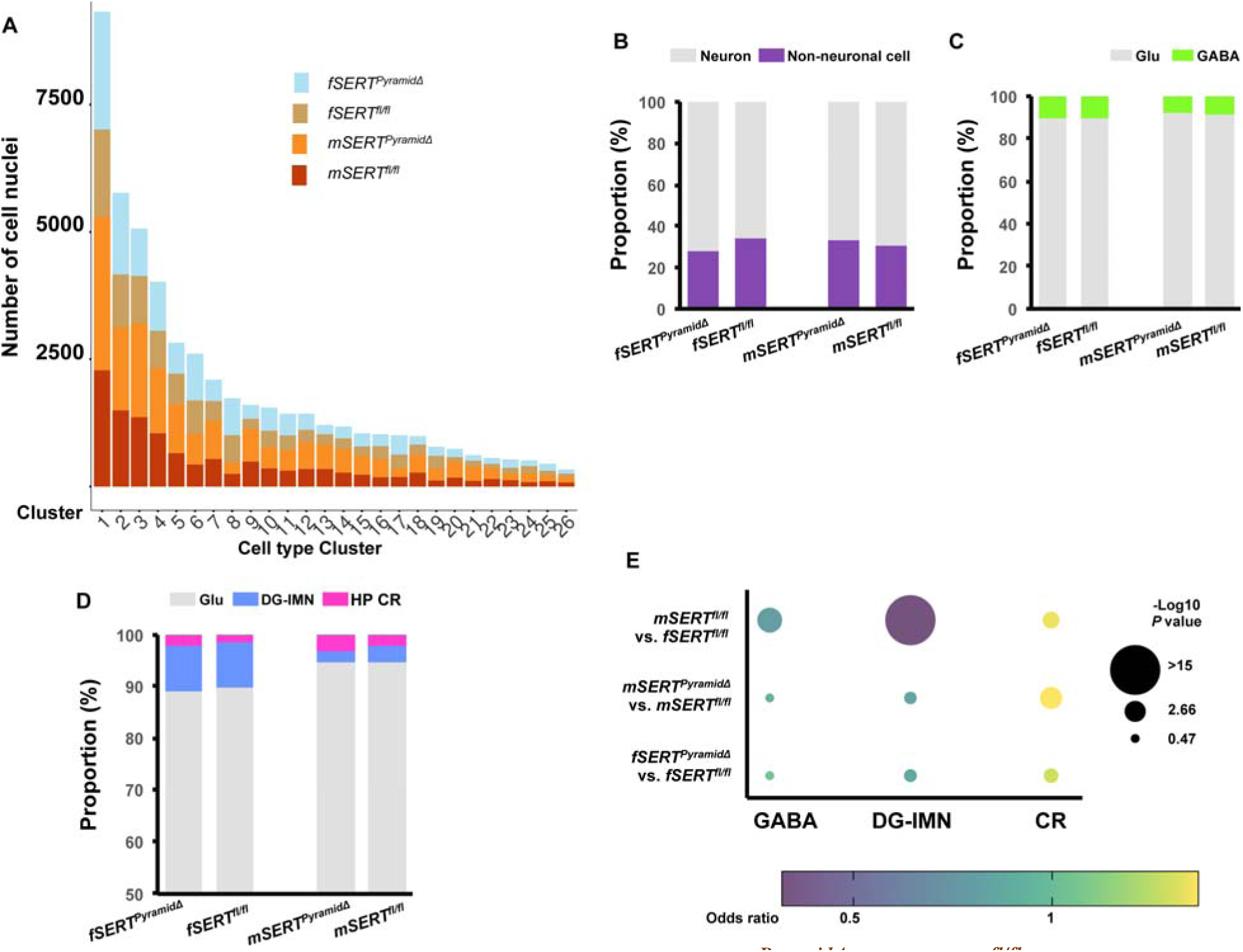
Proportion of transcriptomic clusters in *SERT^PyramidΔ^* and *SERT^fl/fl^* dHPF. **A.** The number of nuclei in individual cell-type clusters in the four samples. **B.** Bar plots showing the proportions of neurons *vs.* non-neuronal cells in the four samples **C.** Bar plots showing the proportions of glutamatergic neurons *vs.* GABAergic neurons in the four samples. **D.** Bar plots showing the relative proportions of glutamatergic neurons, DG-IMNs and HP-CR cells in the four samples. **E.** Dot plots of the odds ratios (OR) of the proportions of GABAergic neurons, DG-IMNs and HP-CR cells in the dHPF transcriptomes for indicated sample pairs. Color and circular size indicate OR of the proportions and null-hypothesis test -Log_10_ *P* value, respectively. OR>1 indicates a higher proportion of the cell types in the first term of the comparison, and OR<1 indicates a lower proportion.

While the cell types were generally balanced among the samples (Figure 3A; Table S1), differences in the proportion of several cell types between the sexes are notable. The proportion of GABAergic neurons was lower in the males (*mSERT^PyramidΔ^* 7.89%, *mSERT^fl/fl^* 8.29%) *vs.* the females (f*SERT^PyramidΔ^* 10.62%, f*SERT^fl/fl^*10.11%) (Figure 3C; Table S1). To evaluate whether the differences were greater than expected by chance, we performed null-hypothesis testing for calculated odds ratio (OR) of the cell proportions between the samples. The proportion of GABAergic neurons was significantly lower in *mSERT^fl/fl^ vs.* f*SERT^fl/fl^* (-Log_10_P = 3.71 Fisher’s exact test), but similar between *mSERT^PyramidΔ^* and *mSERT^fl/fl^* (-Log_10_P = 0.47) and between f*SERT^PyramidΔ^* and f*SERT^fl/fl^* (-Log_10_P = 0.49) (Figure 3E; Table S2).

Likewise, the proportion of DG-IMNs was nearly twice lower in *mSERT^fl/fl^ vs.* f*SERT^fl/fl^* (-Log_10_P >15), but similar comparing *SERT^PyramidΔ^ vs*. sex-matched controls (-Log_10_P =0.94/male, -Log_10_P =1/female) (Figures 3D and 3E; Table S2).

Moreover, the proportion of CR cells was high in *mSERT^fl/fl^ vs.* f*SERT^fl/fl^* (-Log_10_P = 1.68), and became significantly higher in both the *SERT^PyramidΔ^* males (-Log_10_P =2.92) and females (-Log_10_P =1.3) *vs*. sex-matched controls (Figures 3D and 3E; Table S2).

While more samples are needed to strength our results, the findings are interesting. During the first-two postnatal weeks, the numbers of mature GABAergic neurons and DG granule cells arise to build local synaptic circuits (Le Magueresse and Monyer, 2013; Wamsley and Fishell, 2017). In parallel, the vast majority of CR cells are eliminated through programmed cell death (Elorriaga, Pierani et al. 2023). The rates of postnatal neuronal cell generation, differentiation and death differ between the two sexes (Premachandran et al., 2020). Our observations indicate that those fundamental sexually dimorphic processes are preserved in the *SERT^Pyramid^*^Δ^ HPF.

The tight control of CR cell numbers and timing of the death are crucial to HPF functional circuitry (Anstotz et al., 2016; Del Rio et al., 1996; Elorriaga et al., 2023; Mienville and Pesold, 1999). Increased CR cells in postnatal brain have been associated with temporal lobe epilepsy in humans and impaired memory in mice (Blumcke et al., 1999; Elorriaga et al., 2023; Newell et al., 2021). The increased CR cell proportion in the male and female *SERT^PyramidΔ^* dHPF relative to sex-matched controls suggest that CA3 SERT regulates HPF circuits development in the inherent sex context.

### *SERT^PyramidΔ^* alters postnatal gene expression in selective HPF neuronal types

The well-established synaptic connectivity between HPF regions (Figure 4A) (Ding et al., 2020; Lavenex and Banta Lavenex, 2013) permits delineation of cellular and anatomical origin regulated by the transient CA3 SERT expression during behavioral circuits development. Our previous studies observed sex-biased deficits in long-term activity-dependent hippocampal synaptic plasticity and cognitive behaviors in adult *SERT^PyramidΔ^* mice (De Gregorio et al., 2022). SERT-expressing CA3 pyramidal neuron axonal terminals (Schaffer collaterals) innervate the stratum oriens and stratum radiatum of the hippocampal proper (CA1, CA2 and CA3) and the inner-molecular layer of the DG (Figures 1A, 1B and 4A) (De Gregorio et al., 2022). We reasoned that if CA3 SERT influences behavioral circuits through regulating 5-HT at their synaptic partners, we could expect sex-biased gene expression changes enriched in CA1, CA2, CA3 and DG. Alternatively, CA3 SERT may control regional trophic 5-HT, thereby influencing gene expression patterning across the HPF.

**Figure 4.**
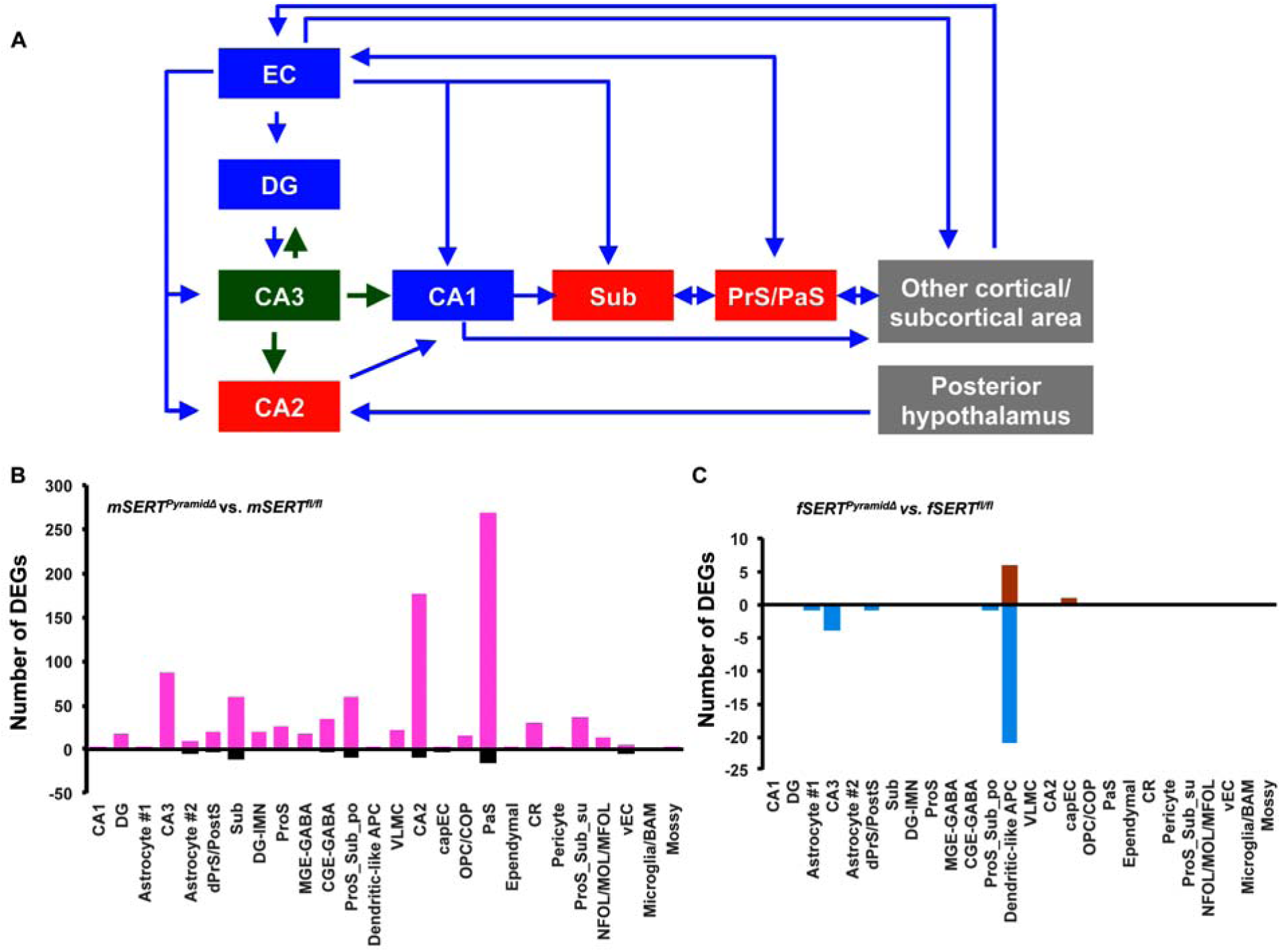
Effects of *SERT^PyramidΔ^* on gene expression in dHPF cell types in males and females. **A.** Schematic of simplified synaptic connectivity of HPF subregions, highlighting CA3 pyramidal neurons do not form synaptic connectivity with subicular regions. **B and C**. Summary of the number of differentially expressed genes (DEGs) in individual transcriptomic cell types in dHPF, comparing m*SERT^PyramidΔ^ vs.* m*SERT^fl/fl^* (**B**), and f*SERT^PyramidΔ^ vs.* f*SERT^fl/fl^* (**C**). Positive and negative numbers indicate upregulated and downregulated genes in *SERT^PyramidΔ^* dHPF, respectively. DEGs were selected based on the statistical thresholds: expression in >10% cells in the cluster, log2FC > 0.25 or < -0.25, FDR <0.05

We performed differential gene expression analysis for every cell cluster, and identified differentially expressed genes (DEGs) at the statistical thresholds of the expression in >10% cells in the cluster, log2-fold change (log2FC) >0.25 or < -0.25, false discovery rate (FDR) < 0.05 comparing sex-matched *SERT^PyramidΔ^ vs*. *SERT^fl/fl^* cells, and additionally log2FC changes in the two sexes are different by 0.2. This analysis identified a total of 1041 DEGs over all the cell types, comprising 707 unique genes with 196 of which altered in multiple cell types (Figures 4B and 4C; Table S3). The majority of the DGEs were in glutamatergic neurons in the *SERT^PyramidΔ^* males, with the greatest number in the CA2 (186 DEGs) and PaS (285 DEGs), sizeable DEGs in the CA3 and Sub, but very few in the CA1 and DG (Figure 4B; Table S3). There was no correlation between the number of DEGs and the population size of the cell type clusters (Figures 3A, 4B and 4C; Tables S1 and S3). These data suggest that SERT expression in the 5-HT-absorbing CA3 pyramidal neurons regulates 5-HT not merely at their synaptic partners, but across the HPF to influence particular aspects of circuits development.

### *SERT^PyramidΔ^* impairs CA2 pyramidal neuron gene expression patterning during social behavioral circuit maturation

The CA2 mediates sociocognitive processes (Diethorn and Gould, 2023; Hitti and Siegelbaum, 2014). CA3 pyramidal neuron Schaffer collaterals innervate CA2 pyramidal neuron dendrites, and CA2 pyramidal neuron soma morphology, dendritic branching and spine distribution/density are sex dimorphic (Bartesaghi and Ravasi, 1999; Dudek et al., 2016). CA2 pyramidal neurons reconfigure the gene expression pattern during functional maturation to elaborate adult cellular properties and synaptic structures (Dudek et al., 2016). In mice, CA2 pyramidal neuron functional remodeling occurs in the first two postnatal weeks, leading to a drastic behavioral switch from the preference for littermates to preference for novel peers around P14 (Diethorn and Gould, 2023; Dudek et al., 2016). Therefore, SERT expression in CA3 5-HT-absorbing pyramidal neurons coincides with the development of adult CA2 pyramidal neuron characteristics and social novelty preference.

To assessing whether the identified CA2 DEGs represent coordinated expression changes of genes involved in discrete cellular functions, we performed Gene Ontology (GO) term enrichment analysis. Of 186 DEGs in the CA2 pyramidal neurons, 177 were upregulated in *SERT^PyramidΔ^* males *vs*. *SERT^fl/fl^* males (Figure 4B; Table S3). Cellular compartment GO enrichment analysis identified 49 upregulated genes in aspects of synaptic structures, including glutamatergic synapse, axon, growth cone, lamellipodium, dendrite and dendritic spine, with some DEGs associated with multiple GO terms (Figures 5A, 5C – 5E; Table S4). In addition, upregulated genes were enriched in cytoplasmic stress granule and endosome (Figures 5A and 5F; Table S4).

**Figure 5.**
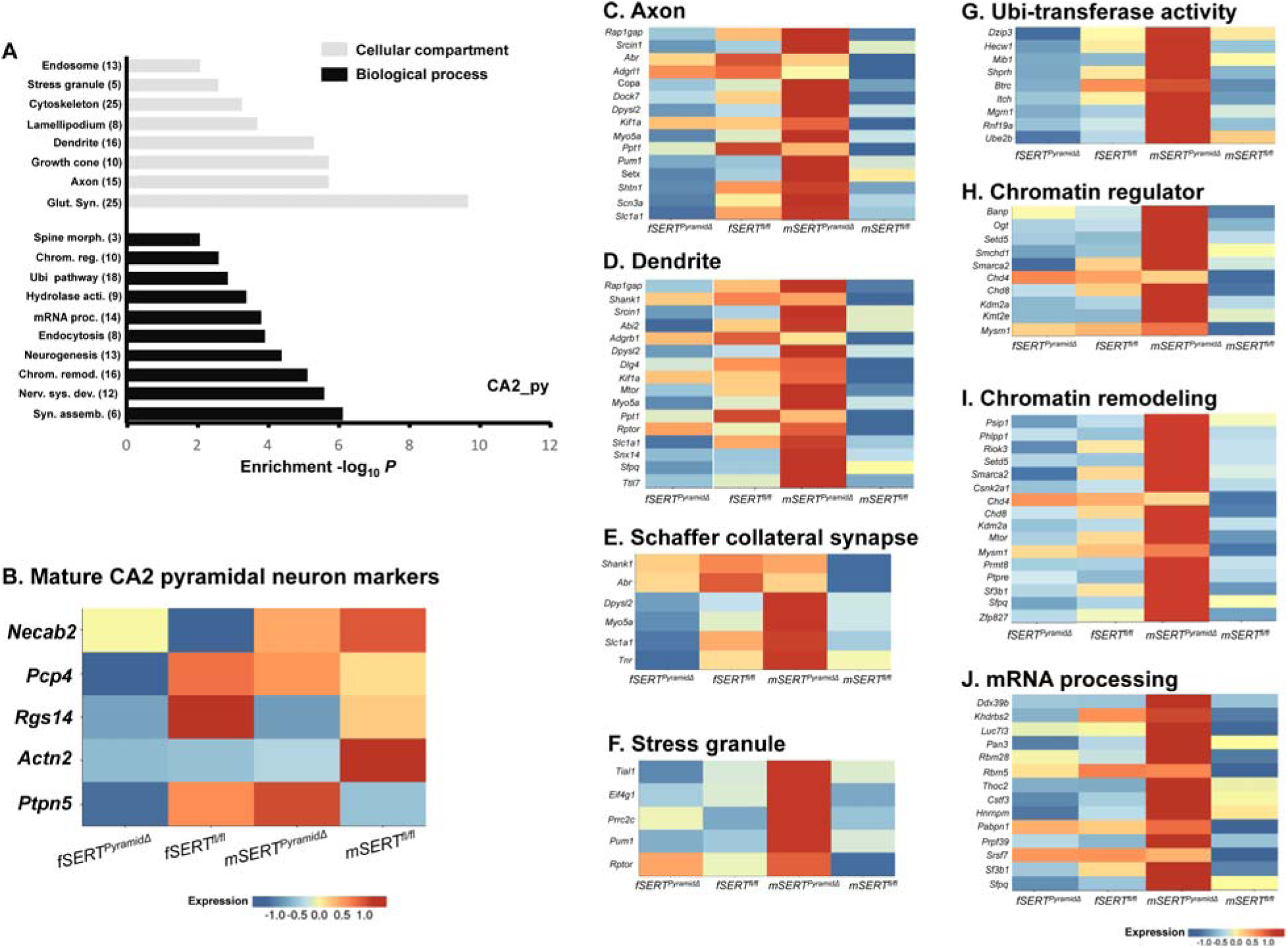
Effects of *SERT^PyramidΔ^* on male CA2 pyramidal neuron gene expression. **A.** Gene Ontology (GO) terms enriched with DEGs in male *SERT^PyramidΔ^* CA2 pyramidal neuron transcriptomes. Grey bars denote GO terms categorized by cellular compartments, and black bars denote biological processes. The number of DEGs matching each category is indicated in parentheses, and the genes in each term are listed in Table S4. **B.** Heatmap showing the expression of gene markers for mature CA2 pyramidal neurons in the CA2 transcriptomes of male and female *SERT^PyramidΔ^* and *SERT^fl/fl^*dHPF. **C. – J.** Heatmaps showing the expression of the DEGs matching indicated GO terms in male *SERT^PyramidΔ^* CA2 pyramidal neuron transcriptomes, relative to that in the other samples. Colors indicate relative cluster mean expression levels.

Aligned with the enrichment of the DEGs in synaptic components, biological process enrichment identified terms of neural development (e.g. synapse assembly, dendritic spine morphogenesis, neurogenesis) (Figure 5A; Table S4). Moreover, the upregulated genes were enriched for chromatin remodeling, mRNA-processing, and Ubi-mediated protein degradation (Figures 5A, 5G – 5J), suggesting that CA3 SERT regulates molecular processes critical for CA2 pyramidal neuron gene expression patterning during the functional maturation.

To study this in more details, we asked whether CA3 SERT expression is essential for CA2 pyramidal neuron functional maturation to occur. Postnatal onset of *Necab2*, *Pcp4, Rgs14, Actn2 and Ptpn5* expression marks mature CA2 pyramidal neurons (Diethorn and Gould, 2023; Dudek et al., 2016). We found all these genes expressed in the male and female *SERT^PyramidΔ^* CA2 pyramidal neurons, albeit some displayed lower levels *vs.* sex-matched *SERT^fl/fl^*CA2 cluster (Figure 5B). This suggests that *SERT^PyramidΔ^* did not obliterate CA2 pyramidal neuron maturation, but altered gene expression patterns critical to the synaptic transmission.

While the DEGs were identified in *SERT^PyramidΔ^* males, there were several notable trends in the expression in the female *SERT^PyramidΔ^* CA2 pyramidal neurons although the changes did not reach our thresholds. Many of the DEGs in GO terms associated with synaptic function tended to change in an opposite direction in f*SERT^PyramidΔ^ vs*. f*SERT^fl/fl^* (Figures 5C – 5E). In contrast, some genes in the stress granules (e.g. *Prrc2c*, essential for stress granule assembly) appeared upregulated in both the *SERT^PyramidΔ^* males and females *vs*. sex-matched controls (Figure 5F). some DEGs in the Ubi-pathway terms also displayed opposite directional changes between *SERT^PyramidΔ^* males and females *vs*. sex-matched *SERT^fl/fl^* controls (Figure 5G). Yet, the DEGs in the biological processes regulating transcriptional plasticity (i.e. chromatin remodeling, chromatin regulators and RNA processing) displayed *SERT^PyramidΔ^* males-specific upregulation (Figures 5H - 5J).

The CA2 pyramidal neuron transcriptomes corroborate bulk RNA-seq of P7 *SERT^PyramidΔ^* HPF (De Gregorio et al., 2022) in that DEGs were preferentially enriched for synaptic function. While the mechanisms underlying CA2 pyramidal neuron functional remodeling are not well understood, male and female social behaviors differ. We observed sex-biased changes in the DEGs associated with synaptic function and male-specific DEGs in chromatin remodeling and RNA-processing in *SERT^PyramidΔ^* CA2 pyramidal neurons at P16, the timing of social novelty emerging. These observations raise the idea that the gene expression patterning during CA2 pyramidal neuron functional maturation, thus the development of the social representation, is regulated by sex-biased mechanisms, which may be modulated by SERT expression in the CA3 5-HT-absorbing pyramidal neurons

### *SERT^PyramidΔ^* impairs gene expression patterning in parasubiculum glutamatergic neurons during spatial circuit development

The subicular complex, located between CA1 and EC, is the output component of HPF signals. It comprises four major subregions ––prosubiculum (ProS), subiculum proper (Sub), presubiculum (dorsal PrS/dPrS is also termed as post-subiculum/PostS) and parasubiculum (PaS), each distinguished by cellular characteristics, long-range inputs, local wiring and projection targets, but not innervated by CA3 pyramidal neurons (Chen et al., 2022; Ding, 2013; Ding et al., 2020; Witter et al., 1989). We identified glutamatergic neuron transcriptomes for each subicular subregion, the ProS and Sub superficial layer (ProS_Sub_su) and deep layer polymorphic cells (ProS_Sub_po) (Figures 2B and 2C; Table S1). Combined, 31% of the total glutamatergic neuron transcriptomes represent the subicular complex. Of the 1041 DEGs identified, 514 were in subicular glutamatergic neurons, with 269 upregulated and 16 downregulated in the male *SERT^PyramidΔ^* PaS *vs.* male control (Figure 4B; Table S3).

The PaS processes spatial information (Dalton and Maguire, 2017; Witter et al., 2014). It forms reciprocal connections relaying information to a suite of cortical/subcortical regions as well as from diverse brain regions into the HPF via medial entorhinal cortex (MEC) (Figure 4A) (Ding, 2013; Witter et al., 2014). Among PaS glutamatergic neurons are cells representing external space –– grid cells, head direction cells and border cells (Boccara et al., 2010; Tang et al., 2016; Witter et al., 2014). In rodents, PaS space cells grow axons, increase branching and innervate targets during the first two postnatal weeks, reaching adult-like functional state at P15, right at the time of eyelid parting and onset of exploratory behavior (Canto et al., 2019; O’Reilly et al., 2013).

Upregulated genes in the male *SERT^PyramidΔ^* PaS glutamatergic neurons were, as in the CA2 pyramidal neurons, enriched for GO terms associated with synaptic transmission, endosome, Ubi-pathway, chromatin regulators/remodeling and RNA processing (Figure 6A; Table S4). Like in the CA2 pyramidal neurons, many DEGs associated with synaptic function and Ubi-pathway displayed opposite trends in female *SERT^PyramidΔ^ vs.* female *SERT^fl/fl^* controls, and the DEGs associated with chromatin plasticity and RNA processing displayed *SERT^PyramidΔ^* male-specific upregulation (Figures 6B - 6E). However, although the GO terms were in common between the CA2 pyramidal neurons and PaS glutamatergic neurons, DEGs in the terms largely differed. In GO terms for chromatin regulator/remodeling, 32 genes were upregulated in the PaS glutamatergic neurons and 20 upregulated in the CA2 pyramidal neurons, only three overlapped (Figure S3A; Table S4). In GO terms for synaptic compartment, 58 genes were upregulated in the PaS and 43 upregulated in the CA2, 14 overlapped (Figure S3B; Table S4). In addition, unlike in the CA2 pyramidal neurons, upregulated genes in the PaS glutamatergic neurons were enriched for transcriptional regulators (33 genes) (Figure 6A; Table S4), suggesting that CA3 SERT may influence PaS glutamatergic neuron functional maturation through global modeling of the gene expression patterns, protein homeostasis, as well as dedicated transcription factors. Together, the data suggest that CA3 SERT expression regulates certain shared cellular processes during functional maturation of CA2 pyramidal neurons and PaS glutamatergic neurons, and specific genes affected by *SERT^PyramidΔ^* may reflect on cell type-specific functional properties.

**Figure 6.**
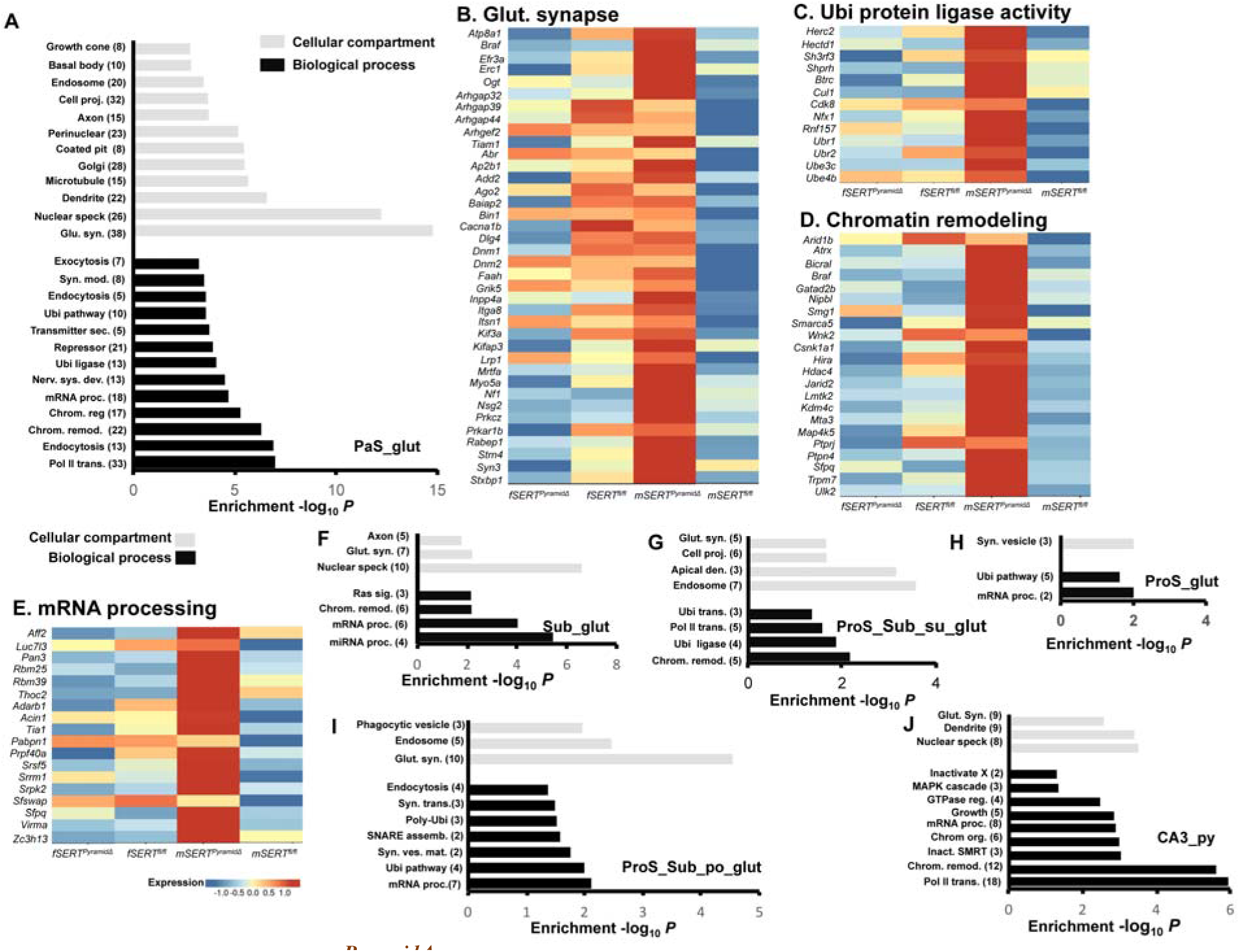
Effects of *SERT^PyramidΔ^* on male PaS glutamatergic neuron gene expression. **A.** Gene Ontology (GO) terms enriched with DEGs in male *SERT^PyramidΔ^* PaS glutamatergic neuron transcriptomes. **B – E**. Heatmaps showing the expression of the DEGs of indicated GO terms in male *SERT^PyramidΔ^* PaS transcriptomes, relative to that in the other samples. **F – J**. GO terms enriched with DEGs in male *SERT^PyramidΔ^* Sub (**F**), ProS_Sub_su (**G**), ProS_glut (**H**), ProS_Sub_po (**I**) glutamatergic neuron transcriptomes, and CA3 pyramidal neuron transcriptomes (**J**). Grey bars denote GO terms categorized by cellular compartments, and black bars denote biological processes. The number of DEGs matching each category is indicated in parentheses, and the genes in each term in corresponding cell types are listed in Table S4. Colors in the heatmaps indicate relative cluster mean expression levels.

Consistently, although the numbers of DEGs in glutamatergic neurons of the other subicular subregions were smaller, *SERT^PyramidΔ^* male-biased upregulated genes were also enriched for GO terms associated with synaptic transmission, chromatin remodeling and RNA processing (Figures 6F - 6I; Table S4). Since subicular glutamatergic neurons are not innervated by CA3 pyramidal neurons, CA3 SERT may regulate trophic 5-HT levels at the extracellular space of the subicular complex to in turn influence the gene expression. The substantially greater number of DEGs in the PaS than the other subicular subregions indicates PaS glutamatergic neuron functional maturation particularly sensitive to trophic 5-HT regulation. As PaS space cells display adult-like function at P15, before the emergence of mature space cells in the EC and place cells in the CA1, the PaS is thought to provide the primitive space representation, setting up developmental trajectories of the spatial circuits (Langston et al., 2010; Witter et al., 2014). Our data suggest that SERT expression in CA3 5-HT-absorbing pyramidal neurons regulates PaS glutamatergic neuron gene expression patterning during the spatial presentation formation.

### *SERT^PyramidΔ^* impairs gene expression patterning in CA3 pyramidal neurons

SERT-expressing CA3 pyramidal neuron axonal projections innervate the stratum oriens and stratum radiatum of CA3 itself, CA2 and CA1, and the inner-molecular layer of the DG both at the ipsilateral HPF and passing the midline via the commissural pathway to the contralateral HPF (Figure 1B) (De Gregorio et al., 2022). Therefore, CA3 SERT is positioned to coordinately regulate 5-HT in the HPF of the two brain hemispheres. Like in the CA2 pyramidal neurons and subicular glutamatergic neurons, *SERT^PyramidΔ^* male-biased upregulation of genes in the CA3 pyramidal neurons were enriched for glutamate synapse, chromatin regulator/remodeling and RNA processing (Figure 6J; Table S4). Again, the DEGs in the terms were largely CA3 specific (Figures S3A and S3B). In addition, *SERT^PyramidΔ^* male-biased upregulated genes in the CA3 pyramidal neurons were enriched for transcriptional regulators and signaling molecules (Figure 6J; Table S4). 5-HT has been shown to control left-right patterning during chick and frog morphogenesis (Fukumoto et al., 2005; Levin et al., 2006). Our observations raise the possibility that regulation of 5-HT by the SERT expression in CA3 pyramidal neurons represents a mechanism orchestrating HPF synaptic circuits development in the two brain hemispheres.

### *SERT^PyramidΔ^* alters ASD-associated gene expression in glutamatergic neurons in specific HPF subregions

Human SERT/*Slc6a4* variants that reduce its expression/function are associated with compulsive behaviors, anxiety disorders and ASD (Abbott et al., 2018; Canli and Lesch, 2007; Caspi et al., 2010; Tordjman et al., 2001). Moreover, early life exposure to SERT inhibitors (i.e. selective 5-HT reuptake inhibitors/SSRIs) increases the risk for ASD (Glover and Clinton, 2016; Harrington et al., 2013; Lugo-Candelas et al., 2018; Olivier et al., 2013). Bulk RNA-seq of P7 HPF showed both male and female *SERT^PyramidΔ^* DEGs highly enriched for ASD-associated genes, and male, not female, *SERT^PyramidΔ^* DEGs enriched for ASD-syndromic genes curated by the SFARI database (De Gregorio et al., 2022). We therefore mapped the SFARI-ASD genes onto our datasets. We found 153 of the 672 (22.8%) male and 10 of the 35 (28.6%) female *SERT^PyramidΔ^* DEGs matched SFARI-ASD genes (Figure 7A; Table S5). Further, male (47 of 672) but not female (2 of 35) DEGs were significantly enriched for SFARI-ASD-syndromic genes (Figure 7A; Table S5).

**Figure 7.**
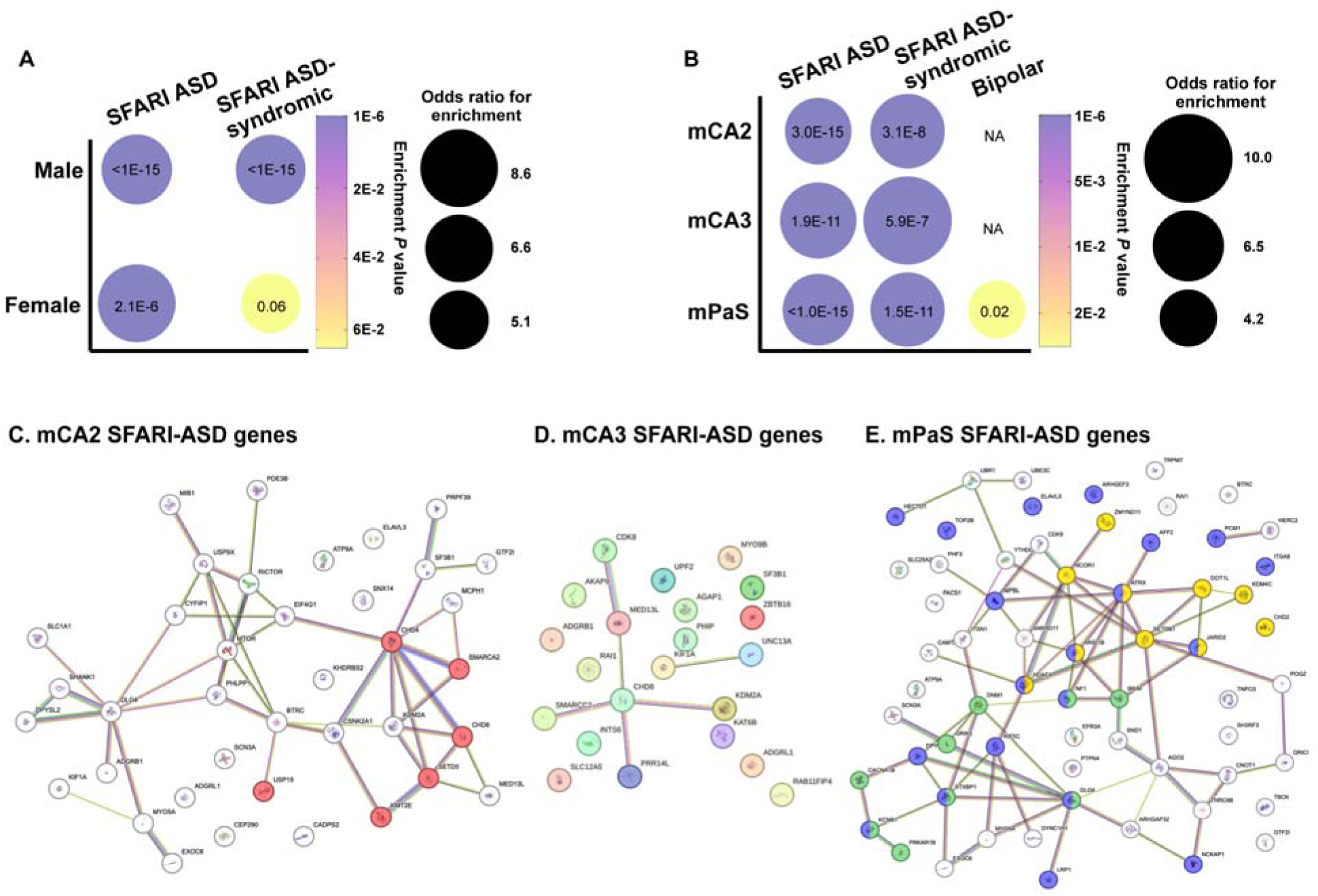
Enrichments of ASD risk genes in DEGs in *SERT^PyramidΔ^* dHPF snRNA transcriptomes. **A.** Enrichments of ASD-associated and ASD-syndromic genes curated by the SFARI database in total DEGs in male and female *SERT^Pyramid^*^Δ^ dHPF. **B.** Enrichments of ASD-associated genes, ASD-syndromic genes and bipolar-associated genes in DEGs in male *SERT^Pyramid^*^Δ^ CA2 and CA3 pyramidal neuron, and PaS glutamatergic neuron transcriptomes. Circle size indicates odd ratio of the fold enrichment and color indicates Fisher’s exact test *P* value as specified in individual circles for the enrichment. The disease genes that matched DEGs in indicated cell populations are listed in Table S5. **C – D.** StringDB network plots showing functional associations of SFARI ASD genes that matched male *SERT^Pyramid^*^Δ^ DEGs in CA2 (**C**), CA3 (**D**) and PaS (**E**). The nodes are genes and the edges represent functional interaction defined by the STRING. ASD genes in the CA2 pyramidal neurons in GO term histone binding (FDR = 0.023) colored in orange, and ASD genes in PaS glutamatergic neurons in GO terms chromatin organization (FDR = 0.026), nervous system development (FDR = 0.022) and modulation of chemical synaptic transmission (FDR = 0.022) in yellow, purple and green, respectively.

Developing glutamatergic neurons are a key point of functional convergence of pleiotropic ASD risk genes (Parikshak et al., 2013; Willsey et al., 2013). To identify ASD genes that are coordinately regulated in specific HPF glutamatergic neuronal types by CA3 SERT during the critical period of circuit development, we mapped SFARI-ASD genes to DEGs in the male *SERT^PyramidΔ^* CA2 pyramidal neurons, CA3 pyramidal neurons and PaS glutamatergic neurons. 20.4%, 25.6% and 22.5% of the DEGs in the CA2, CA3 and PaS matched ASD-associated genes, and 7.5%, 10.5% and 7.4% matched ASD-syndromic genes, respectively (Figure 7B; Table S5). These findings are congruent with the hypothesis that specific groups of ASD-risk genes may form spatiotemporal coregulatory networks influencing distinctive brain developmental processes (Parikshak et al., 2013; Willsey et al., 2013).

To explore spatiotemporal ASD-risk gene networks regulated by CA3 SERT, we examined whether those affected ASD-risk genes converge on particular molecular pathways and cellular processes in CA2 pyramidal neurons, CA3 pyramidal neurons and PaS glutamatergic neurons, using pathway enrichment analyses. We observed neuron-distinct ASD-risk gene functional networks (Figures 7C – 7E). In the CA2, many affected ASD-risk genes intersected in GO term histone binding (FDR = 0.023), including the male-biased ASD-syndromic gene *Chd8* (Figure 7C). This ASD-risk histone modification gene network linked cascades of ASD risk genes implicated in diverse cellular processes (e.g. X-chromosome encoded Ubi-enzyme Usp9, signaling molecules *Mtor*, *Rictor*, postsynaptic glutamate receptors organizer *Dlg4*, high-affinity glutamate transporter *Slc1a1*/EAAT3) as well as the male-biased ASD-risk gene *Shank1* (Figure 7C). In the CA3 by contrast, the affected ASD-risk genes were not enriched for any GO term and did not converge on particular functional pathways, suggesting their engagements into a wide range of CA3 cellular pathways (Figure 7D). In the PaS, however, the affected ASD-risk genes were tightly coalesced in GO terms chromatin organization (FDR = 0.026), nervous system development (FDR = 0.022) and modulation of chemical synaptic transmission (FDR = 0.022) (Figure 7E), linking ASD-risk chromatin plasticity to ASD-risk genes involved in developmental programing of synaptic transmission during PaS spatial representation development.

Dysregulation of 5-HT signaling is implicated in a wide range of behavioral and mood disorders including depression and bipolar disorders and their genetic susceptibilities overlap (Garbarino et al., 2019; Lucki, 1998; State and Sestan, 2012). The *SERT^PyramidΔ^* transcriptomes present an opportunity to explore distinguished cellular and developmental origins as well as shared pathophysiological liabilities of distinct neurological disorders. We tested this idea by matching *SERT^PyramidΔ^* DEGs to bipolar disorder-associated genes (O’Connell et al., 2025). We found no match between the bipolar disorder-associated genes and the DEGs in the *SERT^PyramidΔ^* females, male *SERT^PyramidΔ^* CA2 and CA3 pyramidal neurons, but there were four bipolar disorder-associated genes among the male *SERT^PyramidΔ^* PaS DEGs (Figure 7B; Table S5).

Together, our observations suggest that the perinatal SERT expression in the 5-HT-absorbing pyramidal neurons preferentially regulates the expression of distinct sets of ASD susceptible genes in distinct dHPF glutamatergic neurons during behavioral circuit development. Whereas dHPF is critical to cognitive functions, vHPF is known to modulate emotional states (e.g. stress and anxiety) (Fanselow and Dong, 2010; Kheirbek and Hen, 2011). Thus, it remains plausible that *SERT^PyramidΔ^* affected the expression of bipolar disorder-associated genes in specific neuronal types in the vHPF but not that in the dHPF.

## DISCUSSION

Coherent cognitive behavior requires precise coordination of neural circuits that process related information. Neuronal gene expression patterns and synaptic connectivity are dynamically reconfigured by genetic programing and environmental cues during functional maturation to shape circuit properties (Cadwell et al., 2019; Rakic et al., 2009). Alterations in neural circuit assembly are believed to be a major source of inter-individual variations in cognition and susceptibilities to neuropsychiatric disorders (Bale et al., 2010; Marin, 2016; Silbereis et al., 2016). One unresolved puzzle has been how development of functional related neural circuits is coordinated. This work using single-nucleus transcriptomics of *SERT^PyramidΔ^* dHPF aged P16, along with previous anatomical, electrophysiological and behavioral studies (De Gregorio et al., 2022; Soiza-Reilly et al., 2019), identifies the transient perinatal SERT expression in 5-HT-absorbing pyramidal neurons as a mechanism dedicated to regulate neuronal gene expression patterning during HPF circuit development. Our data provide a basis for linking changes in specific genes in specific cell types in specific HPF regions during the functional maturation to changes in circuit properties and relevant behaviors.

Our results highlight the role of CA3 SERT in coordinating HPF social and spatial representation development. The enrichment in the *SERT^PyramidΔ^* CA2 and PaS of the DEGs involved in chromatin remodeling, RNA-processing, Ubi-pathways and aspects of synaptic transmission suggests that, by controlling trophic 5-HT, the transient SERT expression in those 5-HT-absorbing neurons may provide a means to shape the expression pattern of diverse cellular processes in the time window of social and spatial representation formation. Mapping SFARI-ASD genes to the *SERT^PyramidΔ^* CA2 and PaS transcriptomes revealed alterations of cell-type distinct sets of ASD risk genes, indicating that the cellular processes regulated by CA3 SERT are susceptible to mutations in disparate ASD-risk genes. Our data point to alterations in the co-expression networks during CA2 and PaS functional maturation as an origin of a variety of genetic and environmental disturbances ultimately contributing to shared behavioral phenotypes.

### SERT expression in CA3 pyramidal neurons coordinately regulates social and spatial circuits development

For concerted circuits development, timing is key. This study was motivated by the idea that the exquisite temporal-specific SERT expression in perinatal CA3 pyramidal neurons offers a unique window to reveal certain fundamental principles of functional HPF circuit development. The HPF is a multicomponent brain region mediating declarative learning and relational memory (O’Keefe and Nadel, 1978). Since all the HPF regions receive inputs from raphe serotonergic neurons (Bang et al., 2012; Ding, 2013), we had speculated two plausible outcomes: *SERT^PyramidΔ^* alters gene expression patterning in (1) CA3 pyramidal neuron synaptic partners, i.e. CA3, CA2, CA1 and DG, or (2) all HPF regions due to dysregulated trophic 5-HT.

We unexpectedly observed DEGs preferentially in the CA2 pyramidal neurons and PaS glutamatergic neurons. Since CA3 pyramidal neurons do not innervate the PaS, CA3 SERT likely serves to control extracellular trophic 5-HT levels in the HPF. This assumption is in line with previous findings that 5-HT-absorbing neurons quickly degrade imported 5-HT, and that in *C. elegans* and mice disrupting SERT function in 5-HT-absorbing neurons results in excessive 5-HT signaling across the target region (Chen et al., 2015; Jafari et al., 2011; Persico et al., 2001). As such, our data suggest that CA3 SERT selectively controls 5-HT signaling that regulates CA2 pyramidal neuron and PaS glutamatergic neuron functional maturation.

Why CA2 and PaS? The perception of objects, space and time drives behavior and memories (Squire and Wixted, 2011). CA2 pyramidal neuron synaptic activity controls both social interaction behavior and the passage of time –– the perception of who and when, thus the novelty (Clein and Gould, 2025; Dudek et al., 2016). In rodents, CA2 pyramidal neuron maturation is marked by the emergence of its cellular properties critical for novelty recognition and behavioral switch to favoring novel peers at the end of the second postnatal week (Diethorn and Gould, 2023). Perturbing or delaying CA2 pyramidal neuron maturation permanently impairs social behaviors (Clein and Gould, 2025). Importantly, functional maturation of CA2 pyramidal neurons parallels with the maturation of PaS space cell morphology, synaptic connectivity and electrophysiology (Canto et al., 2019; Witter et al., 2014). Perhaps, the temporal coordination of the gene-expression networks during the CA2 pyramidal neuron and PaS glutamatergic neuron (e.g. space cells) functional maturation coordinates the development of the neural presentations of who, when and where, thereby binding social experiences with emotional valances of places.

The selectivity of *SERT^PyramidΔ^* effects on PaS is noteworthy. Spatial circuits in the HPF involve multiple space cell types: grid cells, head-direction cells and border cells encoding positional metrics, and place cells recording location associated with specific environmental contexts (Witter et al., 2014). PaS, dPrS/PostS and MEC contain grid cells, head-direction cells and border cells and form reciprocal connections; MEC sends efferent to CA1, which contains place cells, thereby binding matric framework to specific contexts (Witter et al., 2014). It has been proposed that PaS and dPrS/PostS form primitive spatial representations, instrumental in building the temporal and spatial code of downstream space cells in the circuits (Canto et al., 2019; Langston et al., 2010; Tang et al., 2016). Indeed, PaS and dPrS/PostS synaptic projections arrive MEC at P4 - P6, becoming fully functional at P11/P12, followed by the emergence of functional grid cells in the MEC at P15/P16, MEC morphological/electrophysiological maturation at P28-P30, and CA1 place cell maturation at P27-P50 (Canto et al., 2019; Langston et al., 2010). Since PaS and dPrS/PostS space cells reach adult-like states before eyelid parting, their spatial representations are thought to be instructed more by genetic programing than environmental experiences (Canto et al., 2019; Langston et al., 2010). The transient CA3 SERT expression may influence the timing and plasticity of the genetic programing of the primitive spatial representations. As PaS and dPrS/PostS connect distinct MEC layers and play distinct biological roles (Canto et al., 2019), *SERT^PyramidΔ^* effects on PaS could reflect on neuron-specific genetic programing.

SERT is also expressed in mPFC L5/6 pyramidal neurons; *SERT^PyramidΔ^* ablates SERT expression in the mPFC pyramidal neurons as well as in the CA3 (De Gregorio et al., 2022). SERT-expressing mPFC pyramidal neurons project to a wide range of cortical and subcortical regions including amygdala and VTA that are critical for cognitive processes (De Gregorio et al., 2022; Soiza-Reilly et al., 2019). Thus, we cannot exclude the possibility that *SERT^PyramidΔ^* impairs mPFC targeting brain regions, which in turn influence CA2 and PaS glutamatergic neuron gene expression. Moreover, SERT is also expressed during this period in thalamocortical projection neurons to define the sensory map in the cortex (Chen et al., 2015; De Gregorio et al., 2020). Thus, another attractive scenario could be that SERT expressing CA3, mPFC and thalamic projection neurons coordinate 5-HT signaling in the development of relevant neural circuits in the HPF, cortex, and subcortical regions that perceive, storage and respond to social encounters and space contexts.

### SERT expression in pyramidal neurons on postnatal HPF gene expression patterning

Analyzing DEGs in male *SERT^PyramidΔ^* dHPF glutamatergic neurons suggest several mechanistic roles of SERT in HPF circuits development. First, global modulation of neuronal plasticity. Across dHPF regions, DEGs fall into two major mechanistic categories: a) regulators of gene expression and protein levels, including chromatin remodeling, RNA-processing and Ubi-dependent protein degradation, and b) aspects of synaptic transmission. Chromatin plasticity is a major mechanism regulating memories (Santoni et al., 2024), and transcriptional and RNA-processing dysregulation underscore neuronal dysfunction seen in ASD (Gandal et al., 2022; Voineagu et al., 2011). Our data add Ubi-dependent protein degradation modulated by CA3 SERT to the plasticity of glutamatergic neuron functional maturation. While the mechanistic categories affected by *SERT^PyramidΔ^* were shared in common between the HPF regions, specific DEGs within the categories and their co-expression networks were neuron-type specific, suggesting SERT, thus 5-HT, regulating functional properties unique to the neuronal types.

Second, preferential effects on glutamatergic neurons. Gennady Buznikov and Jean Lauder have long proposed that 5-HT is an ancient morphogen controlling spatial-temporal organization of ontogenesis (Buznikov et al., 2001). Since glutamatergic neuron projections define the framework of neural circuits, regulation of trophic 5-HT by SERT expression in the 5-HT-absorbing neurons may represent an evolutionary conserved ontogenetic mechanism in shaping functional neural circuit architecture. Indeed, ablating SERT expression in the thalamocortical axons (*SERT^TCAΔ^*) disrupts the sensory map in the cortex (Chen, Ye et al. 2015). *SERT^TCAΔ^* also impairs GABAergic synaptic architecture in the sensory cortex (Chen et al., 2015; De Gregorio et al., 2020). However, we observed small numbers of DEGs in the GABA transcriptomes (Table S3). This could suggest that SERT primarily regulates glutamatergic neuron functional maturation, which in turn shape their partner GABAergic neuron synaptic architecture, thereby coordinating the development of the local glutamatergic-GABAergic circuits with cross-region circuits.

Third, sex-biased *SERT^PyramidΔ^* effects on HPF gene expression. Male and female HPF have distinct patterns of gene expression across development, as a result of chromosomal dosage differences, sex-distinct neuroepigenetic mechanisms and sex hormones (Bundy et al., 2017; Lenz et al., 2012; Premachandran et al., 2020). Thus 5-HT predictably influences gene expression in the inherent sex-biased context. Indeed, our previous studies observed sex differences in the *SERT^PyramidΔ^* effects on P7 HPF bulk transcriptomes, CA1 dendritic architecture, long-term synaptic plasticity and behaviors (De Gregorio et al., 2022). Therefore, we can expect sex-biased DEGs in HPF cell types. Nevertheless, the small number of the DEGs in female *SERT^PyramidΔ^* cell clusters is surprising. The bulk RNA-seq of P7 HPF showed that *SERT^PyramidΔ^* females tend to alter genes involved in translation, while *SERT^PyramidΔ^* males preferentially alter genes critical for transcription (De Gregorio et al., 2022). Because SERT expression in the CA3 terminates by P10, our datasets from P16 HPF reflect on the lasting effects resulting from ablating perinatal CA3 SERT function. Therefore, one explanation could be that the altered translational regulatory mechanisms lead to lasting changes in the synaptic connectivity and activity observed in *SERT^PyramidΔ^* females. An intriguing observation is the trend of opposite changes in the expression levels of the DEGs associated with aspects of synaptic transmission between male and female *SERT^PyramidΔ^* CA2 and PaS *vs.* sex-matched controls. This may be analogous to the male-biased upregulation and male-female opposite changes in gene expression seen in the mouse model for ASD-associated allele of the chromatin-remodeling factor *Chd8* (Jung et al., 2018). It is also worth considering that small changes of multiple components may aggregately impact on biological processes and neural circuitry contributing to certain observed *SERT^PyramidΔ^* female-biased synaptic and behavioral phenotypes.

We acknowledge limitations of our study. Our observations are limited to one developmental time point. Although pooling sex/genotype-matched dHPF from independent cohorts may minimize inter-animal variations, the number of our sequencing samples was small. In particular, we observed very few downregulated DEGs, which could be in part due to smaller magnitude reductions of lower abundance mRNAs not discernible in our single nucleus transcriptomics. Sequencing additional samples will strengthen our findings. Moreover, systematic profiling *SERT^PyramidΔ^* HPF cell types across developmental milestones is necessary to track whether changes in the gene expression during the neuronal maturation result in persistent changes throughout the life, and if they trigger secondary changes over the time, and how these changes influence the perceptions, learning, memories, and behavior.

Our data provide additional evidence that SERT expression in 5-HT-absorbing neurons serves to regulate trophic 5-HT; how trophic 5-HT influences neuronal gene expression, however, is not known. We confirmed all the fourteen 5-HT receptor subtypes expressed in the HPF (Figure S4). We observed most neuronal types co-expressed multiple 5-HT receptor subtypes, particularly *Htr1f, Htr2c, Htr4 a*nd *Htr7*, contrasting to largely absent expression in the non-neuronal types (Figure S4A), very similar to spatially resolved single cell transcriptomics of the 5-HT receptors in the mouse brain (De Filippo and Schmitz, 2024). Like our P7 bulk HPF transcriptomics (De Gregorio et al., 2022), *Htr2c* showed a trend towards a higher expression in *SERT^fl/fl^* females *vs*. *SERT^fl/fl^*males (Figure S4B). Interestingly, *Htr4* displayed a trend of opposite directional changes between male and female *SERT^PyramidΔ^ vs.* sex-matched controls in the CA2 pyramidal neurons, CA3 pyramidal neurons and PaS glutamatergic neurons (Figures S4C – S4E). 5-HT receptor subtypes may form heteromultimer complexes coupled to distinct intracellular signaling pathways (Berumen et al., 2012; Wirth et al., 2017). Therefore, trophic 5-HT may influence social and spatial representation development via 5-HT receptor downstream signaling pathways. Another intriguing possibility is that trophic 5-HT may directly influence gene expression, through a recently described 5-HT-modification of chromatin by covalently attached to the histone protein H3 at glutamine 5 (histone H3 serotonylation) (Farrelly et al., 2019). Trophic 5-HT may readily enter diverse cell types, diffuse between the cytoplasm and nucleus and interact with “intracellular targets” (Buznikov et al., 2001; Colgan et al., 2009). Possibly, the dynamic chromatin structural plasticity in CA2 pyramidal neurons and PaS glutamatergic neurons during the functional maturation render them particularly sensitive to the changes in trophic 5-HT signaling. Further studies are needed to delineate neuron-specific trophic 5-HT signaling pathways. Nonetheless, the findings presented here provide a working framework for mechanistic investigation into genetic programing of social and spatial circuit assembly in shaping cognition and behaviors.

## MATERIALS and METHODS

### Mice

Animal use and procedures were approved by the institutional animal care and use committee at the Albert Einstein College of Medicine. Generation of *SERT^fl/fl^* mice has been previously described (Chen et al., 2015). *SERT^Pyramid^*^Δ^ mice were generated by crossing *SERT^fl/fl^;Emx1-Cre* (*Emx1-Cre*/Jackson Laboratories 00562, (Gorski et al., 2002)) females and *SERT^fl/fl^* males, and sex-matched *SERT^fl/fl^* littermates were the controls. Figures 1A and 1B were adapted from our published work, and experimental methods utilized in these studies were described (De Gregorio et al., 2022).

### Dorsal HPF dissection

Male and female *SERT^Pyramid^*^Δ^ and sex-matched *SERT^fl/fl^* littermates aged P16 were used in this study. The date of birth of the pups was designated as P0. The pups were genotyped to assess the utility of the litters, ensuring a balanced distribution of genotypes and sexes. 5 pairs of male and 6 pairs of female *SERT^Pyramid^*^Δ^ and *SERT^fl/fl^* mice from multiple cohorts were used. At P16, the mice were anesthetized with an intraperitoneal injection of avertin (400 mg/kg) before being sacrificed by decapitation. The brains were removed, the HPF were isolated free from the EC and surrounding brain tissues using a spatula, and the dorsal portion of the HPF was separated from the ventral portion with a scalpel and flash-frozen in a bath of isopentane on dry ice. dHPFs from the same genotype and same sex were pooled into a single cryovial and stored in liquid nitrogen until nuclei isolation.

### Single-nucleus isolation

The procedures for isolating cell nuclei from frozen dHPF were developed by modification of 10x Genomics “Demonstrated Protocol CG000167 Revision A” and published works (Krienen et al., 2020; Zhou et al., 2020). Specifically, the frozen tissue was thawed on ice for 1 minute, and 500 µL of prechilled Lysis Buffer (10 mM Tris-HCl, 10 mM NaCl, 3 mM MgCl2, 0.025% Nonidet P40 Substitute) was added. The tissue was homogenized using a pestle, gently moving it up and down without twisting. The homogenate was transferred into 5 mL of prechilled Lysis Buffer in a multiwell plate, pre-coated with 1% UltraPure BSA (Thermo Fisher Scientific, AM2616), on ice, and pipetted up and down 20 times using a P1000 pipette. The sample was incubated on ice for 10 minutes, with gentle pipetting every 2.5 minutes to ensure thorough lysis.

The homogenized solution was loaded into a cold 10 mL syringe fitted with a 25-gauge cold needle and expelled into a second well of the multiwell plate. This process was repeated, with the sample transferred to a third well. The solution was then filtered through a 30 µm MACS filter (Miltenyi Biotec, MACS SmartStrainers, #130-098-458) to remove debris, and the filtrate was collected in a prechilled 15 mL Falcon tube pre-coated with 1% BSA. The sample was centrifuged at 500 × g for 5 minutes at 4°C.

After centrifugation, the supernatant was carefully removed without disturbing the nuclei pellet. The pellet was resuspended in 5 mL of prechilled Nuclei Wash and Resuspension Buffer (2.0% BSA, 0.2 U/µL RNase Inhibitor - Takara #2313B - in 1xPBS) and mixed gently by pipetting 20 times with a regular-bore P1000 pipette. The sample was filtered again through a 30 µm MACS filter, and the filtrate was collected in another prechilled BSA-coated 15 mL Falcon tube. This process, including centrifugation and resuspension, was repeated once more. The final pellet was resuspended in 500 µL of chilled Nuclei Wash and Resuspension Buffer and mixed by pipetting 8–10 times. This suspension was combined with 900 µL of a 1.8 M sucrose solution (1.8 M sucrose, 10 mM Tris-HCl, 3 mM MgAc2, 1 mM DTT) and mixed gently. The resulting 1.4 mL solution was layered into a BSA-coated 2 mL Eppendorf tube over 500 µL of a 1.8 M sucrose cushion. The sample was centrifuged at 13,000 × g for 45 minutes at 4°C to separate nuclei from myelin debris. After centrifugation, the supernatant was carefully removed without disturbing the nuclei pellet. The pellet was resuspended in 200 µL of chilled Nuclei Wash and Resuspension Buffer and mixed gently by pipetting 8–10 times. The nuclei suspension was filtered through a 40 µm FlowMi Cell Strainer (SP Bel-Art, #H13680-0040), and the filtrate was collected in a cold BSA-coated 1.5 mL Eppendorf tube. The nuclei were kept on ice and promptly transferred to the Einstein College of Medicine’s Genomics Core for quality assessment, counting, and library preparation.

### Sequencing, data processing and QC

Nuclei quality was assessed, and the concentration determined using a Neubauer Hemocytometer visualized on an Invitrogen™ EVOS™ XL Core microscope. On average, the yield for the four samples was 380,000 nuclei in 200 µL (approximately 1,900 nuclei/µL). From each sample, approximately 16,000 nuclei were loaded targeting a recovery of 10,000 nuclei for single-nuclei RNA library preparation, using the 10x Genomics Chromium Next GEM Single Cell 3’ Reagent Kits v3.1 - Dual Index, following manufacturer’s instructions. Single-nuclei libraries from individual samples were checked for quality and insert size using an Agilent Bioanalyzer with the High Sensitivity kit. The libraries were pooled and sequenced on an Illumina HiSeq 4000 sequencing system using a paired-end 2×150 bp configuration, targeting 350M reads per library to achieve an average of 20,000 reads/nucleus.

### Processing of single nucleus sequencing data

The snRNA-seq reads were aligned and processed by the CellRanger (10X Genomics, version 6.1.1), with the key metrics summarized in Figure S1.

### Integration and comparison of single nucleus sequencing data

The snRNA-seq gene expression matrices from CellRanger were analyzed by the scDAPP pipeline, including QC of individual samples, integration of the four samples, clustering of the integrated data, and differential expression analysis of each of the cell clusters. The non-default parameters included selection of nuclei with 1,000 to 45,000 UMIs, >200 genes and <5% mitochondrial reads, and removal of doublets. The four samples were integrated with the RPCI algorithm (Liu et al., 2021), and 30 principal components were used for Louvain clustering and UMAP visualization. Differential gene expression between *SERT^Pyramid^*^Δ^ and *SERT^fl/fl^* were performed for male and female data separately, using all the nuclei in individual clusters and the MAST algorithm, which considers both bimodal expression distribution and stochastic dropout. DEGs were selected based on the thresholds of expression in >10% cells in the cluster, log2FC >0.25 or < -0.25, and false discovery rate (FDR) < 0.05. By this algorithm and the strict criteria, only few genes were significantly changed in both males and females. We then selected genes with their log2FC changes in the two sexes differed by at least 0.2 for further analysis of sex-biased expression (Table S3). The integrated data were prepared by ShinyCell (Ouyang et al., 2021) for public accessing.

### Cluster odds ratio analysis

To assess whether the differences in the proportion of cell-type clusters between the sexes and genotypes were greater than expected by chance, we performed null hypothesis testing for calculated cluster odds ratio (OR). OR = (a/c)/(b/d), with a and b being the numbers of cluster cells in two comparing samples, and c and d being the numbers of cells in the reference class in the respective samples. The reference cell class for GABAergic neurons was the total neurons minus GABAergic neurons. DG granule cells and glutamatergic neurons were the references for DG-IMN and CR cells, respectively. OR>1 indicates a greater proportion of a cell type in sample a *vs.* sample b, and OR<1 indicates a smaller proportion of a cell type in sample a. OR and null-hypothesis testing *P* values were plotted in the graphs and listed in Table S2.

### GO enrichment analysis

GO term enrichment of the DEGs in individual cell-type clusters and gene hits in each GO term were determined through DAVID functional annotation analyses (https://davidbioinformatics.nih.gov) (Huang et al., 2009; Sherman et al., 2022). Terms for GO cellular compartment and biological processes are reported. An adjusted *P* value <0.05 was used to determine significant terms. Only significant terms are shown in the Figures, -log_10_ *P* value represents GO significance score for each term and specific genes in the terms are listed in Table S4.

### Disease-associated gene enrichment analysis

To assess enrichment of ASD-associated genes in specific cell types of *SERT^Pyramid^*^Δ^ dHPF, we identified overlapping genes between the DEGs in individual cell-types and ASD genes as well as genes classified as syndromic ASD genes curated in the SFARI Gene Autism Database (gene.sfari.org, 10-21-2025 release). Some genes (primarily human- and primate-specific genes) in the SFARI database were not expressed in our datasets, leaving 1,173 ASD-associated and 297 ASD-syndromic genes used in our analyses. To assess the enrichment of bipolar-associated genes, we used gene lists from (O’Connell et al., 2025). The odds ratio for the fold enrichment was calculated as OR= (k/n)/(K/N), with k being the number of DEGs in the disease gene set, n being the number of the DEGs not in the disease gene set; K being the number of non-DEGs in the disease gene set, and N being the number of non-DEGs not in the disease gene sets. The significance of the fold enrichment was calculated by Fisher’s exact test.

To assess functional intersections between SFARI-ASD genes that were dysregulated in *SERT^Pyramid^*^Δ^ HPF particular cell types, the DEGs in the male CA2 pyramidal neurons, CA3 pyramidal neurons and PaS glutamatergic neurons matched SFARI-ASD genes were uploaded to the STRING database. In the interaction network, the nodes are genes and the edges represent functional interaction defined by the STRING, which contains both literature support and prediction. The nodes are further colored by the selected GO terms.

### Statistical analyses

For the cell proportion analyses and disease-associated gene enrichment analyses a two-sided Fisher’s exact test was performed using GraphPad Prism (GraphPad software, v10). Differences were considered statistically significant at *P* < 0.05.

## Acknowledgements

We thank David Reynolds and the Genomics Core Facility at the Albert Einstein College of Medicine for quality assessment of the nuclei and library preparation. This work was supported by NIHR01MH105389 (JYS).

## Author contributions

RDG and JYS conceived of the project, designed and performed the experiments and wrote the manuscript. WC, MAA, RDG and JYS analyzed the snRNA-seq data under DZ supervision. All authors reviewed and edited the manuscript.

## Competing interest

The authors declare no competing interest.

## Notes

https://scviewer.shinyapps.io/hippocampus_sertKO

### Competing Interest Statement

The authors have declared no competing interest.

