## Supplemental Figures for "Hippocampus single-nucleus transcriptomics reveals coordinated regulation of social and spatial representation development by perinatal SERT expression in CA3 pyramidal neurons"

**A**

| | *fSERT*$^{Pyramid\Delta}$ | *fSERT*$^{fl/fl}$ | *mSERT*$^{Pyramid\Delta}$ | *mSERT*$^{fl/fl}$ |
|---|---|---|---|---|
| n_nuclei | 12,863 | 10,351 | 16,525 | 12,308 |
| Total_n_reads | 467,696,017 | 431,896,045 | 437,551,136 | 398,395,477 |
| Median UMI counts per nucleus | 6,604 | 6,490 | 5,374 | 6,506 |
| Median genes per nucleus | 2,467 | 2,466 | 2,231 | 2,511 |
| Total genes detected | 26,224 | 26,158 | 26,378 | 26,054 |

**B**

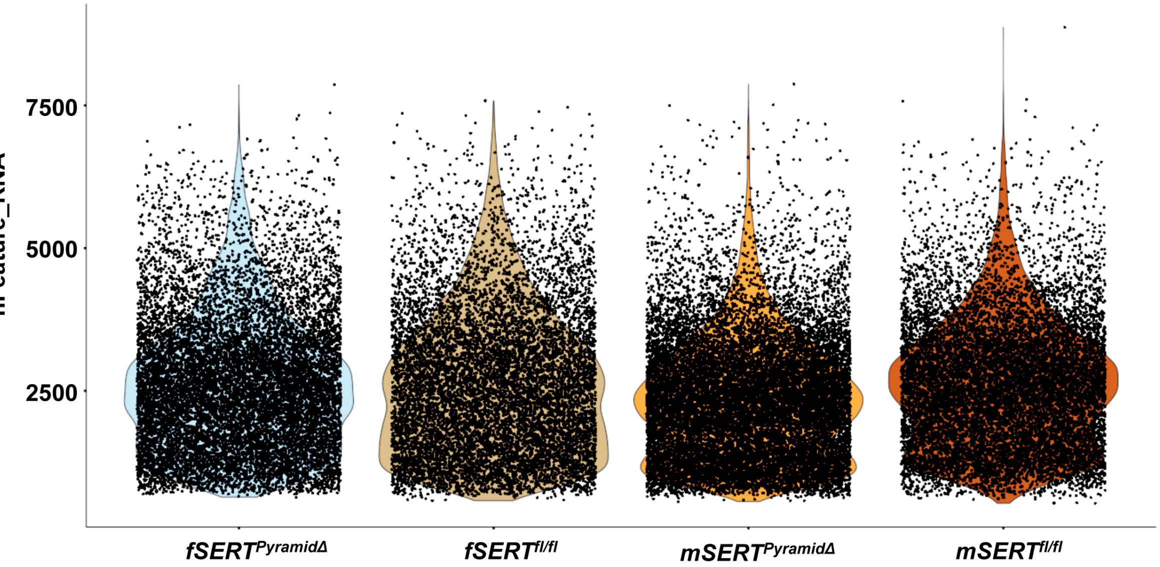


**Figure S1. Key metrics from snRNA-seq of male and female *SERT*$^{Pyramid\Delta}$ and *SERT*$^{fl/fl}$ dHPF**

**A**. Total number of nuclei, sequencing reads, UMI counts per nucleus, median genes per nucleus and genes detected from snRNA-seq of indicated dHPF samples.

**B**. Scatter plots showing the number of genes detected per nucleus (nFeature_RNA) in indicated dHPF samples.

Figure S2

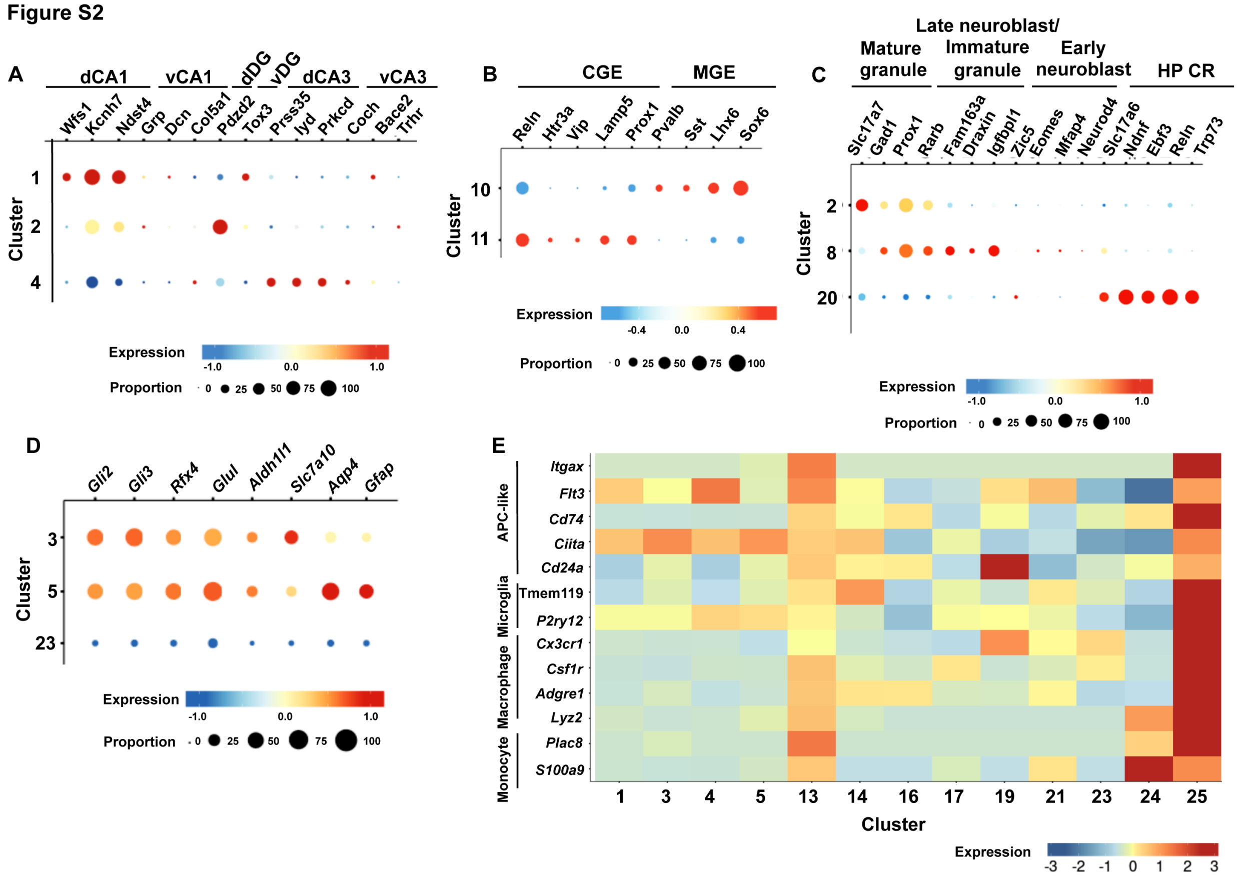


**Figure S2. Selectivity of cell-type markers expressed in transcriptomic cell clusters from snRNA-Seq of P16 dHPF**

**A**. Dot plots showing the expression of marker genes for dorsal and ventral CA1 pyramidal neurons, CA3 pyramidal neurons and DG granule cells in indicated transcriptomic cell clusters.

**B**. Dot plots showing the expression of marker genes for CGE- and MGE-GABAergic neurons in indicated cell clusters.

**C**. Dot plots showing the expression of marker genes for mature DG-granule cells, DG-IMNs at early and late developmental stages, and HP CR cells in indicated cell clusters.

**D**. Dot plots showing the expression of marker genes for astrocytes in indicated cell clusters.

**E.** Heatmap showing the expression of gene markers for various classes of immune cells in indicated clusters.

Dot size and color indicate the proportion of expressing cells and average expression level in each cluster, respectively.

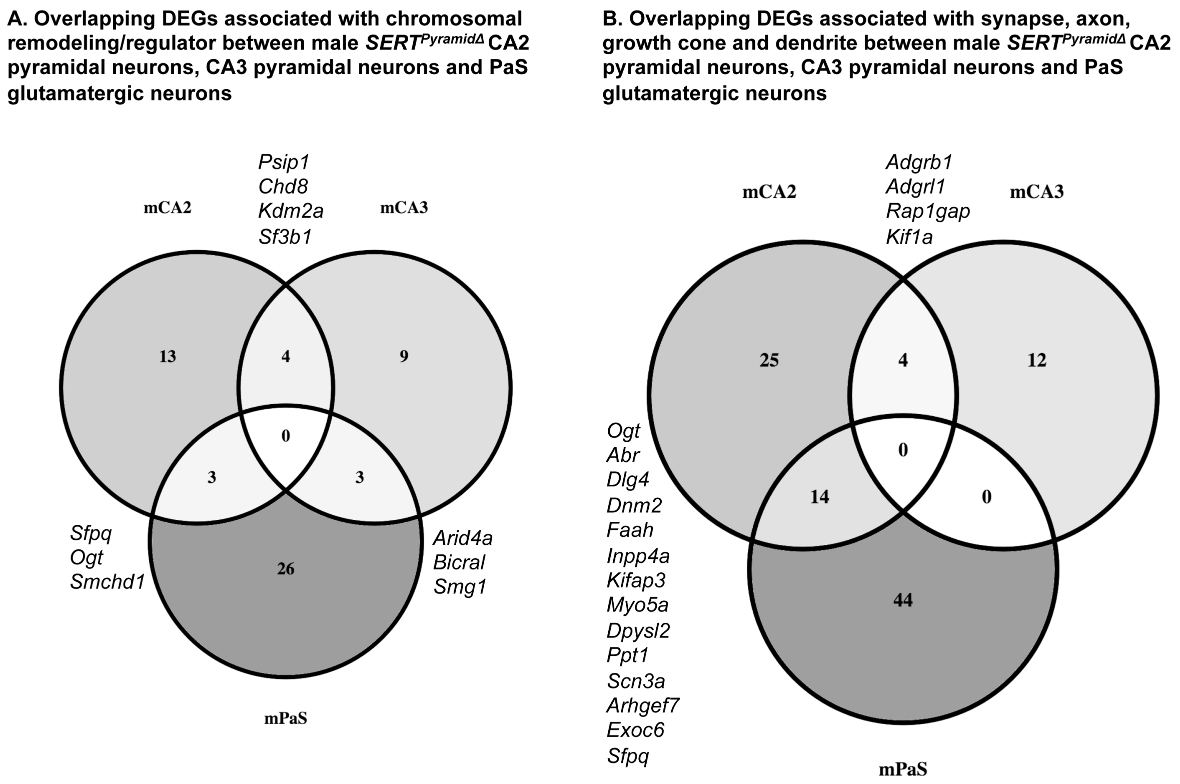


**Figure S3. Shared DEGs in GO terms enriched in CA2, CA3 and PaS.**

**A**. Venn diagram displaying overlap between DEGs in GO terms associated with chromatin remodeling and regulators in male *SERT$^{Pyramid\Delta}$* CA2 pyramidal neuron, CA3 pyramidal neuron and PaS glutamatergic neuron transcriptomes.

**B**. Venn diagram displaying overlap between DEGs in GO terms associated with synaptic compartment in male *SERT$^{Pyramid\Delta}$* CA2 pyramidal neuron, CA3 pyramidal neuron and PaS glutamatergic neuron transcriptomes.

The numbers indicate overlapping and non-overlapping DEGs for each cell cluster, and the overlapping genes are shown next to the two comparing clusters.

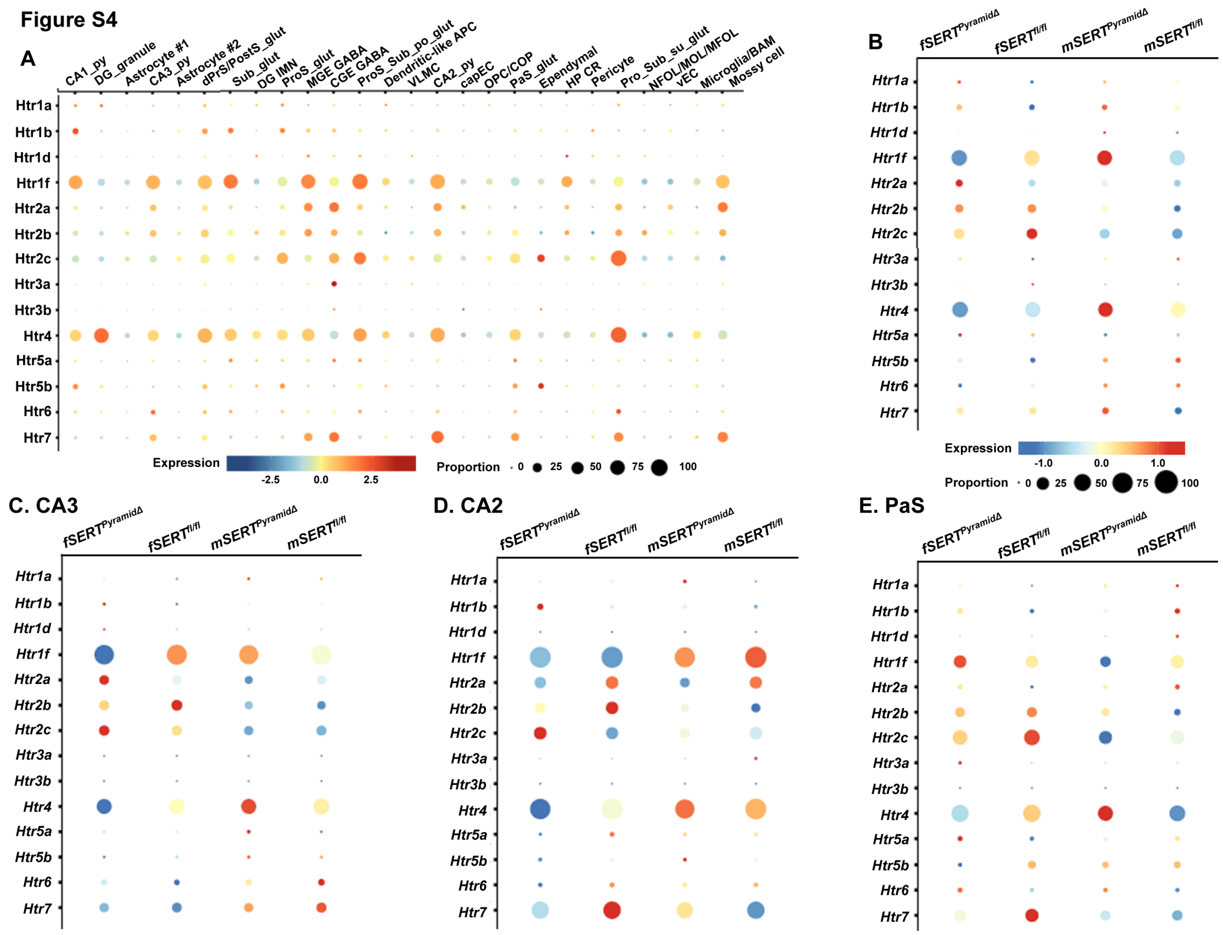


**Figure S4. Expression of fifteen 5-HT receptor subtypes in P16 dHPF cell types**

**A**. Dot plots showing average expression of the 5-HT receptor subtypes by cell types.

**B**. Dot plots showing average expression of the 5-HT receptor subtypes in all transcriptomes in male and female $SERT^{Pyramid\Delta}$ and $SERT^{fl/fl}$ dHPF.

**C – E.** Dot plots showing average expression of the 5-HT receptor subtypes in male and female $SERT^{Pyramid\Delta}$ and $SERT^{fl/fl}$ transcriptomes of CA3 pyramidal neurons **(C)**, CA2 pyramidal neurons (**D**) and PaS glutamatergic neurons (**E**).

Dot size and color indicate the proportion of expressing cells and average expression level in indicated transcriptomes, respectively.
